# Repurposing Melatonin in dual-mode for Wilson disease therapy as a Copper Chelator and an antioxidant agent

**DOI:** 10.1101/2025.08.30.673002

**Authors:** Raviranjan Pandey, Arpan Narayan Roy, Sandip Sarkar, Rakiba Rohman, Kaustav Chakraborty, Rupa Bargakshatriya, Sanjana Pandey, Pruthwiraj, Debosmita Bhattacharya, Saurav Kumar, Saptarshi Maji, Aidan T. Pezacki, Sumit K Pramanik, Christopher J Chang, Ashima Bhattacharjee, Neelanjana Sengupta, Amitava Das, Arnab Gupta

## Abstract

Loss-of-function mutations in copper-ATPase ATP7B underlie Wilson disease (WD), a disorder characterized by hepatic copper accumulation and severe hepato-neuropathology. Existing chelation therapeutics remove excess copper but lack intrinsic antioxidant capacity and frequently cause systemic toxicity. Here we evaluate melatonin, an FDA-approved indoleamine with antioxidant and putative metal-chelating activity, as a candidate therapeutic for WD. In ATP7B⁻^/^⁻ hepatocytes, melatonin restored copper-induced reactive oxygen species (ROS) to basal levels, reduced apoptosis twofold, and attenuated Nrf2 nuclear translocation leading to reduction of Hemoxygenase-1 abundance. Live-cell ratiometric analysis of GSSG/GSH using GRX1-roGFP2 expressed in melatonin-treated ATP7B⁻^/^⁻ hepatocytes revealed a significant reduction in intensity-ratio, indicating an effective mitigation of copper-induced glutathione oxidation. Isothermal calorimetric titration revealed moderate Cu^2+^ affinity (K ^ITC^=4.54 × 10³ M⁻¹), rationalized by MD-simulations showing an interaction energy of 18.5 × 10⁻³ kcal·mol⁻¹ via amide–Cu²⁺ coordination. *In-cellulo* studies also revealed that copper-induced vesicularized ATP7B reinstates to Golgi in melatonin-treated hepatocytes. *In vivo*, melatonin treatment reduced copper-induced oxidative stress in zebrafish embryos and lowered copper burden in *Caenorhabditis elegans* WD model. Our studies revealed that encapsulation of melatonin within an engineered polymeric nanocapsules having dithiol linkers, susceptible to cleavage by GSH, extended melatonin’s circulatory half-life ten-fold and enhanced its ROS-scavenging efficacy three-fold relative to free melatonin. This work introduces a unique dual-function therapeutic strategy that integrates antioxidant activity with copper chelation, simultaneously addressing copper overload and redox imbalance. Repurposing melatonin, with its established clinical safety, offers rapid and cost-effective translational pathway toward WD-therapy while providing a generalizable platform for redox- and metal-associated disorders.

## Introduction

Copper(Cu) is an essential micronutrient for all eukaryotes as it plays a crucial role in numerous biological processes, including mitochondrial respiration, iron metabolism, pathogen neutralization, and antioxidant defence.^1^,^2^ As a redox-active metal, copper serves as a critical cofactor for key enzymes such as cytochrome c oxidase, superoxide dismutase, and ceruloplasmin, which play vital roles in cellular energy production and oxidative stress management.^3^ Specifically, cytochrome c oxidase is integral to ATP production within the mitochondrial electron transport chain, superoxide dismutase mitigates oxidative damage by neutralizing reactive oxygen species (ROS), and ceruloplasmin is essential for iron metabolism and transport.^4^ However, the redox activity that renders copper biologically indispensable also contributes to its potential toxicity when present in excess. In such conditions, copper can catalyse Fenton-like reactions, generating highly reactive hydroxyl radicals that induce lipid peroxidation, protein oxidation, and DNA damage. ^5^ This oxidative stress leads to pathophysiological consequences such as excessive autophagy, endoplasmic reticulum (ER) stress, and cell death. ^6-8^ as the accumulation of copper leads to cellular dysfunction, copper homeostasis needs to be tightly regulated. ^9^

The dual nature of copper is particularly evident in Wilson disease (WD), a genetic disorder caused by mutations in the *ATP7B* gene, which encodes a copper-transporting ATPase responsible for biliary copper excretion. ^10^ In Wilson disease, defective copper excretion results in a toxic accumulation of copper, primarily in the liver and brain in the later stages of the disease. Due to the accumulation of excess copper, redox balance is disrupted in cells, leading to oxidative stress, hepatic dysfunction, fibrosis, and liver failure if untreated. ^11^ However, currently available drugs lack free radical scavenging properties and do not directly overcome the oxidative stress and cellular damage caused by copper overload. This underscores the urgent need for novel therapeutic strategies that regulate copper levels and mitigate oxidative injury. Current therapeutic strategies for Wilson disease focus on copper chelation, e.g., D-penicillamine and Trientine, or zinc supplementation to reduce copper levels by upregulating metallothionein (MT) or inhibiting its absorption.^12^ However, available drugs are often associated with significant side effects, including neurological deterioration, haematological abnormalities, and gastrointestinal disturbances, along with limited efficacy and frequent patient compliance issues.^13^ Specifically, the most commonly administered drug, D-penicillamine, causes worsening of neurological symptoms upon prolonged usage.^14^ Similarly, another potent copper chelator, Trientine, exhibits side effects of sideroblastic anaemia and neurological deterioration.^15^ A table of presently used drugs against WD and their side effects has been tabulated in **Table S1**. Such mild to severe side effects of present WD therapies call for the immediate development of an effective and safer treatment molecule.

Melatonin, an endogenous hormone primarily known for regulating circadian rhythm, can also act as an antioxidant with cytoprotective properties.^16^ Emerging evidence suggests that melatonin exerts protective effects against oxidative stress by scavenging free radicals and modulating key antioxidant pathways, including the nuclear factor erythroid 2-related factor 2 (Nrf2) pathway, which upregulates antioxidant enzymes such as heme oxygenase-1 (HO-1).^17^ Additionally, melatonin has been hypothesized to possess metal-chelating ability, suggesting a dual mechanism of action against copper toxicity.^18-20^ Despite the versatile role of melatonin in the physiological system, its clinical application is limited by its short half-life, rapid metabolism, and potential side effects at high doses. This necessitates the development of advanced delivery systems to enhance its therapeutic efficacy and safety, as well as to enable slow and sustained drug release. ^21^

In this study, we investigated the cytotoxic effects of excess copper on the human hepatic cell line HepG2 and ATP7B knock-out HepG2 (cellular model), as well as organismal models of Wilson disease (*cua-1^-/+^ Caenorhabditis elegans)*, which mimic the hepatic copper accumulation observed in patients. Using a combination of cellular, biophysical, and computational approaches, we explored the interplay between copper toxicity, oxidative stress, and antioxidant and chelating properties of melatonin. To further enhance melatonin’s therapeutic efficacy, we developed a novel polymeric nanocapsule delivery system for stimuli-responsive release under oxidative stress conditions. Our findings establish melatonin’s potential as a multifaceted therapeutic agent for copper overload disorders, capable of both chelating excess copper and mitigating oxidative stress through intrinsic antioxidant activity, especially in Wilson disease and other oxidative stress-related pathologies. We demonstrate for the first time the copper-chelating ability of melatonin using multiple complementary biophysical and cellular approaches, establishing a novel dual-function therapeutic platform for Wilson disease that uniquely integrates copper chelation with antioxidant activity in a stimuli-responsive design—addressing both metal overload and its downstream oxidative consequences, which current therapies fail to achieve. This integrated approach represents a paradigm shift from conventional monofunctional chelation therapy to a next-generation, multifunctional therapeutic platform with broad implications for treating Wilson disease and other redox-driven pathologies.

## Results and Discussion

### Excess copper induces oxidative stress in hepatocytes and decreases cell viability

We investigated the effect of excess copper (Cu) on cell viability and oxidative stress. We observed that when a human hepatic cell line (HepG2) is treated with excess Cu, its viability decreases concentration-dependently **(Fig. 1A)**, which signifies the cytotoxic effect of excess copper on wild-type(wt) HepG2 cells, consistent with known mechanisms of copper toxicity^22^. To examine the underlying cause of the decrease in cell viability, we tested the intracellular oxidative stress level using the fluorescence CellROX assay. CellROX is a cell-permeable fluorogenic probe that is brightly fluorescent upon oxidation by ROS, allowing for quantitative measurement of oxidative stress. Oxidative stress was recorded to be four-fold higher in copper-treated HepG2 cells over untreated ones **(Fig. 1B).** This elevated oxidative stress might be the key contributing factor to decreased cell viability, as ROS are well-established mediators of cellular damage. To further delineate the copper toxicity in the hepatocyte and mimic Wilson disease (WD) conditions, we generated ATP7B knock-out (ATP7B^-/-^) HepG2 using CRISPR-Cas9. Successful knockout of ATP7B was confirmed by immunoblotting **(Fig. 1C)** and immunofluorescence **(Fig. 1D).** Copper level was four-fold higher in ATP7B^-/-^ HepG2 cells than wild-type cells **(Fig. S1A)**, confirming the essential role of ATP7B in hepatic copper export^23^, and consequentially oxidative stress was also elevated **(Fig. S1B)**. In response to copper-induced stress, cells increased autophagy, which is supported by the lipidation of LC3-I to LC3-II **(Fig. S1C)**. This lipidation is a signature event indicating the initiation of the autophagy process^7^. Higher copper accumulation in the KO cell triggered autophagy and apoptosis pathways as measured by cleavage of Caspase-3 and lipidation of LC3I. The cleavage of Caspase-3 serves as a hallmark of apoptosis **(Fig. S1D)**^24^. Copper-induced toxicity was severely elevated in ATP7B^-/-^ cells as the IC_50_ for copper decreased to 569 µM **(Fig. 1E)**, while for wild-type HepG2 cells, the IC_50_ was 964 µM **(Fig. 1A)**. Similar to wild-type cells, oxidative stress was elevated ∼5-fold in KO cells upon copper treatment **(Fig. 1F)**. The stark differences in cell viability and stress markers reestablish the importance of ATP7B in protecting hepatocytes from copper toxicity.

**Figure 1.**
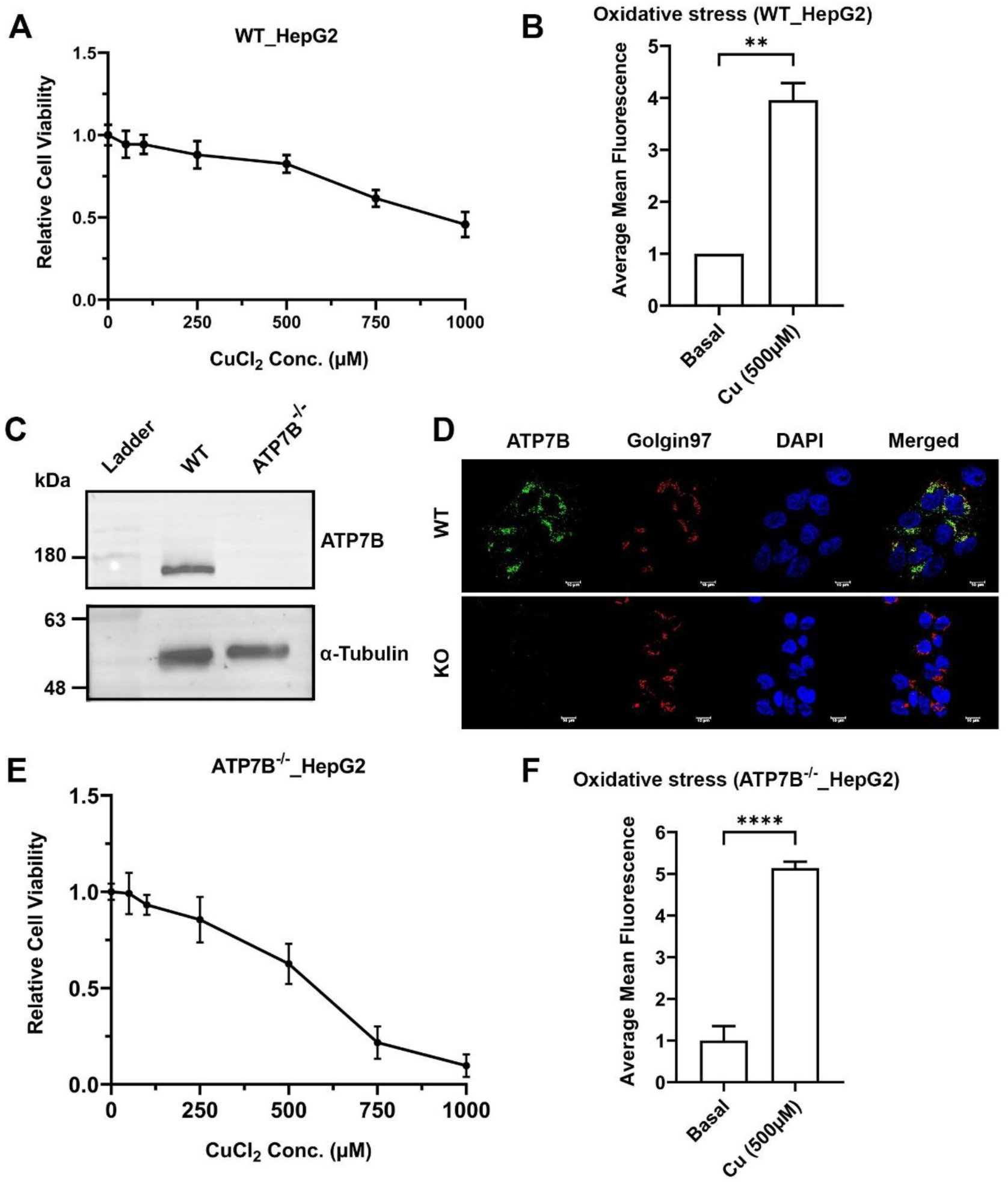
Excess copper induces oxidative stress and cytotoxicity in HepG2 cells, exacerbated in ATP7B-deficient cells. (A) Cell viability of wild-type HepG2 cells treated with increasing concentrations of CuCl2 for 24 h, measured by MTT assay. A dose-dependent decrease in viability was observed. (B) Quantification of intracellular ROS using CellROX Green fluorescence in HepG2 cells treated with 500 µM Cu for 24 h. ROS levels increase ∼4-fold compared to untreated cells (p < 0.01, unpaired t-test, n = 3). (C) Immunoblot confirming successful knockout of ATP7B in HepG2 cells using CRISPR-Cas9. (D) Immunofluorescence analysis validating the loss of ATP7B protein in knockout (KO) cells. (E) Copper cytotoxicity curves of ATP7B^-/-^ HepG2 cells. IC₅₀ for Cu decreased significantly in KO cells (569 µM) versus wild-type (964 µM) (p < 0.01, n = 3), indicating increased sensitivity. (F) ROS levels in ATP7B–/– vs. HepG2 cells after 500 µM Cu treatment. KO cells show ∼5-fold increase in ROS. (p < 0.001, unpaired t-test, n = 3). Data are presented as mean ± SD from three independent experiments. Statistical significance was determined using an unpaired t-test. *p < 0.05; **p < 0.01; \*\**p < 0.001*.

In summary, dose-dependent cytotoxic effects of excess copper on HepG2 cells were observed, and these may be primarily mediated through elevated oxidative stress. The absence of ATP7B significantly aggravates these effects by impairing copper export and further increasing oxidative stress. Thus, a therapeutic candidate with antioxidants and copper-chelating properties may offer a better alternative to reduce copper-induced toxicity and restore cellular copper homeostasis in Wilson disease^25^.

### Melatonin mitigates copper-induced cytotoxicity and oxidative stress in a cellular model of Wilson disease

Melatonin, an endogenous hormone primarily known for regulating circadian rhythm, is also a US FDA-approved dietary supplement. Melatonin is effective in reducing oxidative stress under a large number of circumstances, which is achieved via a variety of means: direct detoxification of reactive oxygen species and indirectly by stimulating antioxidant enzymes while suppressing the activity of pro-oxidant enzymes^26^. It has been hypothesized as a potent therapeutic molecule for the therapy of Wilson disease^27^. We tested the efficacy of melatonin as a therapeutic molecule in conditions of copper accumulation. Melatonin was found to be non-toxic up to 1 mM for at least 24 hours in both wild-type and ATP7B^-/-^ HepG2 cells (**Fig. S2A, S2B**). To test if melatonin could be repurposed as a drug for the therapy of WD, we evaluated its protective effects. Our study reveals that melatonin improves cell viability after copper treatment as the IC50 for copper increases from 526 µM to 4961 µM after melatonin treatment in ATP7B^-/-^ HepG2 (**Fig. 2A**). A similar protective effect was observed in wt HepG2 cells, as the IC50 for copper increased from 964 µM to 14 mM (**Fig. S2C**). This significant improvement in cell viability indicates that melatonin can alleviate copper-induced cytotoxicity in hepatic cells. To understand how melatonin improves cell viability, we measured Cu-induced oxidative stress levels using CellROX after melatonin treatment. In WT HepG2 cells, melatonin treatment significantly reduced Cu-induced oxidative stress to basal levels after 500 µM melatonin treatment (**Fig. 2B**). We also observed a significant decrease in oxidative stress in ATP7B^-/-^ HepG2 (**Fig. 2C**). The fluorescence intensities were quantified in a large cohort of cells (**Fig. 2D,2E**). To further validate our finding, we utilized flow cytometry, which confirmed a significant increase (∼3-fold in wt HepG2 and 4-fold in ATP7B^-/-^ HepG2 cells) in mean fluorescence intensity in Cu-treated cells; this was greatly reduced after melatonin treatment **(Fig. 2F,2G**). To directly assess the antioxidant capacity of melatonin, we performed a DPPH (2,2-diphenyl-1-picrylhydrazyl) free radical scavenging assay. Melatonin demonstrated significant dose-dependent scavenging activity (**Fig. S2D**), confirming its intrinsic, direct antioxidant potential^28^. We have also checked the influence of melatonin on Cu^2+^-mediated hydroxyl radical (•OH) generation, which is one of the main free radicals produced during many cellular processes through a Fenton-like reaction. Using electron paramagnetic resonance (EPR) spectroscopy with DMPO spin-trapping, we observed a strong signal for •OH in the control reaction (Cu^2^⁺ + H₂O₂). The addition of melatonin resulted in a sharp decrease of the EPR signal, indicating significant inhibition of hydroxyl radical formation (**Fig. 2H**). This suggests melatonin directly interferes with the copper-driven Fenton reaction, preventing the generation of the hydroxy radical inside the cellular environment.^29^ These observations demonstrate that melatonin is a promising agent for counteracting copper toxicity as it increases cell viability and reduces copper-induced oxidative stress in wt and ATP7B^-/-^ hepatocytes, supporting its potential therapeutic application ^30^.

**Figure 2.**
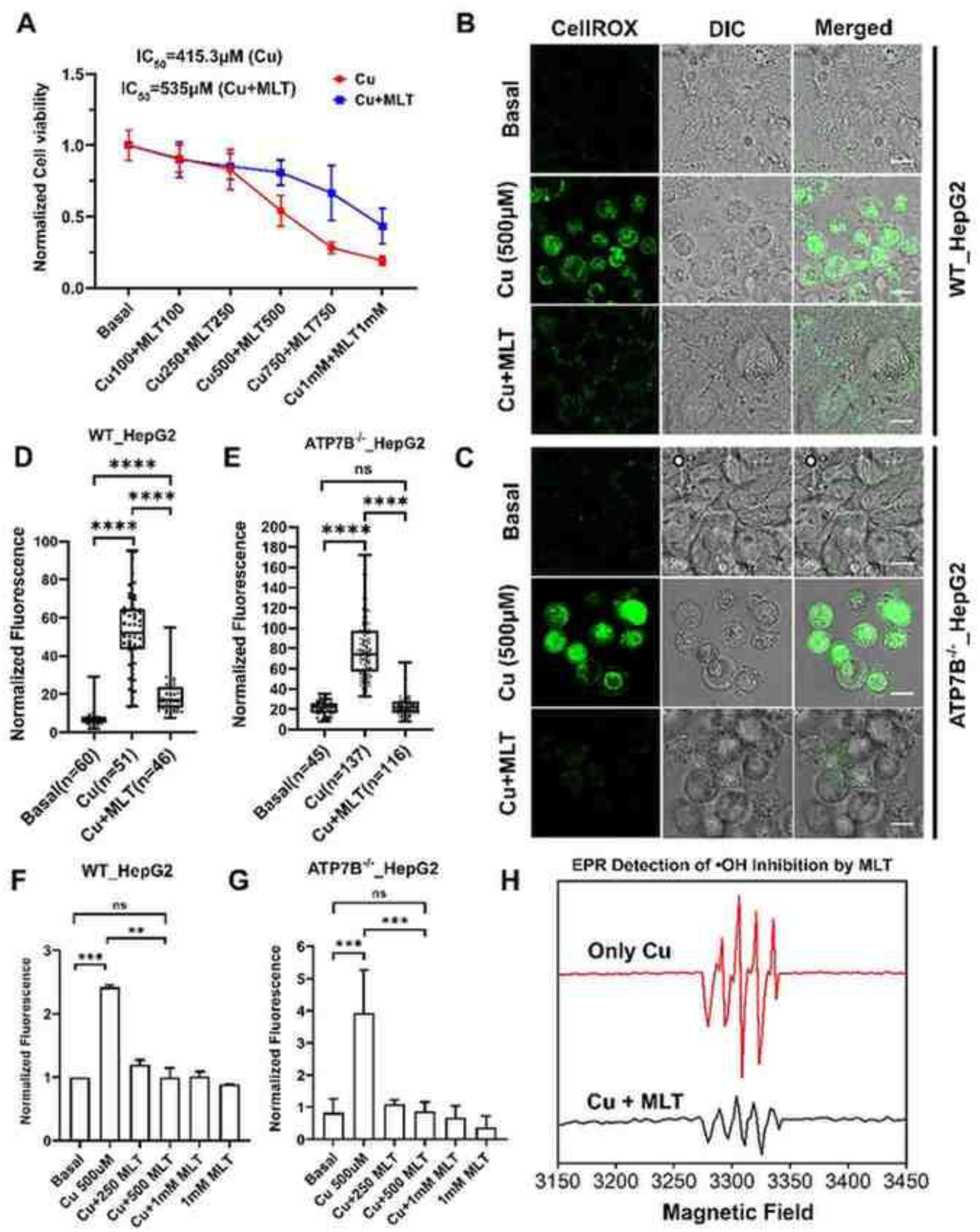
Melatonin mitigates copper-induced cytotoxicity and oxidative stress in wild-type and ATP7B–/– HepG2 cells. (A) Dose–response curve for copper toxicity in ATP7B^-/-^ HepG2 cells with or without melatonin pre-treatment (500 µM, 24 h). IC₅₀ increases from 526 µM (Cu only) to 4961 µM (Cu + melatonin), indicating enhanced cell survival. (B) and (C) ROS levels in wild-type (WT) and ATP7B^-/-^ HepG2 cells after 500 µM Cu treatment with or without melatonin (500 µM), assessed using CellROX Green fluorescence. Melatonin reduces Cu-induced oxidative stress to near-basal levels (*n* = 3). (D) and (E) Quantification of CellROX Green fluorescence intensity in WT HepG2 ATP7B–/– HepG2 cells from three independent experiments. Cu significantly increases ROS levels, which were reduced upon melatonin treatment ( *n* = 3). (F) and (G) Flow cytometric analysis of ROS levels in WT and ATP7B–/– HepG2 cells stained with CellROX Green. Cu treatment increases mean fluorescence intensity (MFI) ∼3-fold in WT HepG2 and ∼4-fold increase in ATP7B–/– HepG2 cells, which was significantly reduced by melatonin (*n*=3). (H) Electron paramagnetic resonance (EPR) spectra using DMPO as a spin-trap to detect hydroxyl radicals (•OH) generated by the Cu^2^⁺ + H₂O₂ system. A strong DMPO–OH signal is detected in the Cu–H₂O₂ mixture, which is markedly attenuated by melatonin, indicating inhibition of hydroxyl radical formation. All data are shown as mean ± SD from at least three independent experiments. Statistical comparisons were made using an unpaired t-test. *p < 0.05; **p < 0.01; \*\**p < 0.001*.

### Spectroscopic and *in silico* studies demonstrate melatonin and copper binding is energetically favourable

Based on a detailed examination of its chemical structure, which features potential N, O-donor atoms, we hypothesized that melatonin can act as a chelator for copper. To probe this, ^1^H NMR titrations were performed with Cu^2+^. The NMR spectra exhibited two significant changes, paramagnetic-induced downfield shifts and peak broadening, and the most prominent effects were observed for the protons adjacent to the amide group **(Fig. 3A, 3B, S3A).** ^31^ Similar shifts were observed upon addition of Cu^+^; this can be rationalized by the facile oxidation of Cu^+^ to Cu^2+^ under aerobic conditions.^32^ The observed paramagnetic shifts and line broadening strongly indicate coordination of Cu^2^⁺-ions through AmideN1–Cu^2^⁺/ Cu^+^ coordination.

**Figure 3.**
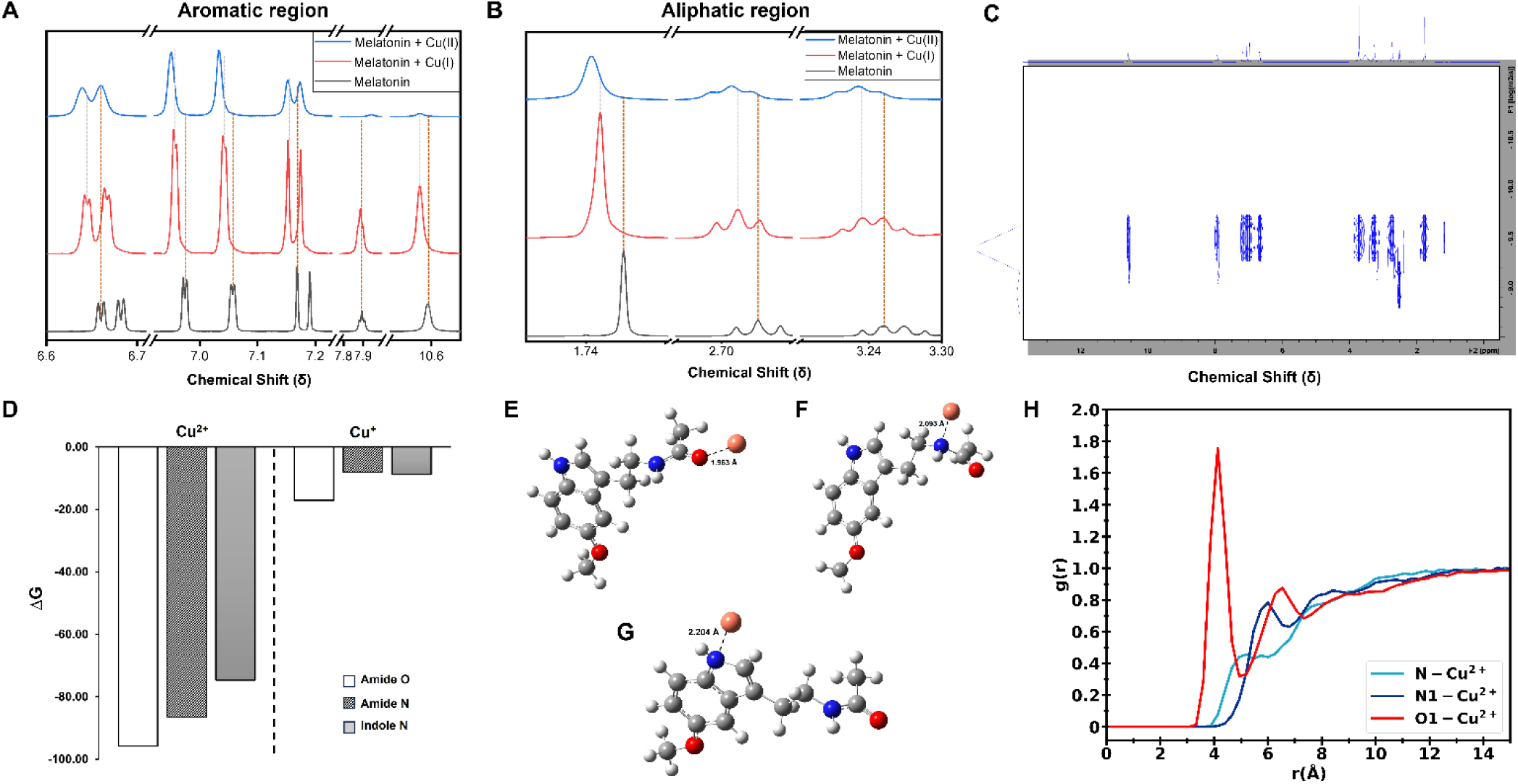
Melatonin preferentially binds Cu^2^⁺ through amidic nitrogen forming a stable coordination complex: evidence from NMR, DOSY, MD, and DFT studies. (A) and (B) Representative ^1H NMR spectra of melatonin showing downfield shifts and disappearance of the amide proton signal upon Cu^2^⁺ binding, suggesting direct coordination via the amide nitrogen (C) DOSY NMR showing a shift in the diffusion coefficient of melatonin from 2.044 × 10⁻^10^ m^2^/s to 2.184 × 10⁻^10^ m^2^/s after Cu^2^⁺ complexation, indicating an altered hydrodynamic radius. (D) Reaction free energies (ΔG, kcal/mol) for Cu^2^⁺–melatonin complexes in aqueous medium *via* the direct chelation mechanism (DCM), computed at the M06-2X/6-311++G(d,p) level. (E, F, G) represent optimized geometries with coordination through the amide O, amide N, and indole N, respectively. Distances between the potential chelation sites [O (*red*) and N (*blue*)] and the Cu atom (*brown*) are indicated with dotted lines. (H) Radial distribution function (RDF) plots of Cu^2^⁺ around the indole nitrogen (N), amide nitrogen (N1) and amide oxygen (O1) of melatonin, extracted from MD simulations.

To further validate complex formation, DOSY NMR experiments were conducted to monitor changes in hydrodynamic volume. Diffusion coefficient of melatonin changes from 2.044 × 10⁻^10^ m^2^/s (Free melatonin) to 2.184 × 10⁻^10^ m^2^/s (Melatonin-copper complex) after Cu^2+^ addition **(Fig. 3C, S3B, S3C, S3D)**. This decrease in diffusion rate indicates a change in hydrodynamic volume, providing evidence for forming a stable coordination complex^33^. Similar changes were observed through ESI–MS analysis, which displayed a distinct mass signal with the characteristic isotope pattern for a copper-containing species at *m/z* 382, corresponding to the [Cu₂(melatonin-H)] ⁺ complex **(Fig. S3E)**. Collectively, these spectroscopic findings identified the primary coordination sites. This guided our subsequent molecular dynamics (MD) simulations and density functional theory (DFT) calculations, which were undertaken to elucidate the precise geometry and energetics of copper coordination. The mechanism of copper binding by melatonin has been investigated first under the DFT framework. Two plausible pathways were considered: the direct chelation mechanism (DCM) and the deprotonation-assisted chelation mechanism (DACM), with the amide and indole groups as potential chelating groups (details are in **S3F**). Three likely chelation sites were examined within these groups, viz., amide O, amide N, and indole N. The calculated free energies (ΔG) for the melatonin-copper complexation, in aqueous medium, revealed exergonic reactions with both Cu^2+^ and Cu^+^, following the DCM pathway (**Table S3F-1**). These results clearly indicated the preference of melatonin to form complexes through the amide group over the indole. Among the three reactive sites, the lowest energy requirement for complex formation was observed for the amide O, making it the most favourable chelation site **(Fig. 3D)**. The reaction feasibility order for Cu^2+^ was obtained as amide O > amide N > indole N. In contrast, for Cu^+^, the order was amide O > indole N ≈ amide N (optimized geometries are presented in **Fig. 3E-3G, and S3F-2, A-C**). In the case of Cu^2+^ complexes, natural population analysis (NPA) indicated the reduction of Cu^2+^ to Cu^+^ upon complexation with melatonin, consistent with previous experimental observations^35^. The Second-order perturbation theory analysis within the NBO framework showed significant donor-acceptor interactions, with electron density delocalization from the lone pairs of the amide nitrogen into the vacant orbitals of Cu. The associated E^2^ energies suggested the monodentate coordination mode of melatonin in these complexes (**Table S3F-2**). In contrast, the DACM pathway, involving bidentate coordination through the amide N and O, and monodentate through indole N, resulted in endergonic reactions of melatonin with both Cu^2+^ and Cu^+^ ions, suggesting lower favourability of chelation of copper ions via this mechanism (**Fig S3F-2, D-G, Table S3F-1 and S3F-2**). Nonetheless, a previous study indicates this type of mechanism (Coupled-deprotonation-chelation mechanism) to be a predominant route for hydrated Cu^2+^ binding^36^ under physiological conditions. We further performed MD simulations to explore the binding behaviour of putative chelation sites of melatonin with copper ions in aqueous media at room temperature. The radial distribution functions g(r) (**Fig. 3H, S3G**) of Cu^2^⁺ around melatonin’s amide N (N_1_), and amide oxygen (O_1_) show sharp peaks at 6.00 Å and 4.13 Å, respectively. Indole nitrogen (N) has a broader peak at 5.20 Å. We observe the sharpest peak corresponding to the amide oxygen (O1), consistent with the DFT estimates.

Our combined spectroscopic and computational studies reveal that melatonin preferably coordinates with Cu ions through its amide group, forming a stable complex in aqueous solution. These findings consistently underscore melatonin’s chelation behaviour under physiological conditions.

### Melatonin sequesters copper *in vitro* and *in cellulo*

Our study has established that melatonin can act as an antioxidant, which aligns well with existing knowledge.^37^ However, conclusive evidence that illustrates copper chelation by melatonin is largely lacking. Fluorescence spectroscopy was performed to determine the binding constant of melatonin and Cu^2+^ ions using a Horiba Fluoromax spectrofluorometer. The intrinsic fluorescence of melatonin (30 μM) was recorded, followed by incremental additions of Cu^2^⁺. A gradual quenching of fluorescence intensity was observed with increasing Cu^2^⁺ concentration, indicating complex formation **(Fig. 4A)**^38^. Linear fitting of the data yielded a binding constant of approximately 1.65 × 10^3^ M⁻^1^ **(Fig. 4B)**. These findings were further supported by isothermal titration calorimetry (ITC), which revealed a binding constant of approximately 4.54 × 10^3^ M⁻^1^, with a binding stoichiometry (n) of 1.59, suggesting the involvement of more than one binding site **(Fig 4C** and **4D)^39^**. Thermodynamic analysis yielded values of ΔG ≈ –4.99 kcal/mol and TΔS ≈ –26.6 kcal/mol, indicating a spontaneous and enthalpy-driven binding process. The ITC data suggest that both nitrogen atoms of melatonin are involved in coordination with Cu^2^⁺, which is further corroborated by in silico molecular docking studies. The negative ΔG value confirms the thermodynamic favourability of copper binding to melatonin. We tested the specificity of the binding of melatonin to copper. Towards that, we measured fluorescence quenching after adding other divalent metal ions, Ca^2+^, Mg^2+^, Mn^2+^, Co^2+^, Zn^2+,^ and Ni^2+^. We did not observe any fluorescence drop for these ions **(Fig. S4 A-F).** This finding also illustrates that essential and non-essential ions in cells will not be sequestered by melatonin and interfere with their physiological roles. We tested whether melatonin can chelate copper in the cellular milieu. Under elevated copper conditions, ATP7B localizes in the trans-Golgi network (TGN) and transports copper in the TGN lumen. Copper treatment triggers vesicularization of ATP7B from the TGN. When we treated the cells with vesicularized ATP7B with melatonin, ATP7B relocalizes to the TGN, an observation reminiscent of cells under copper chelator treatment **(Fig. 4E,4F)**. The finding indicates that melatonin can act as a copper chelator *in cellulo* that induces recycling of ATP7B from copper-induced vesicles to the TGN. ^40^ We also verified the melatonin Cu binding property using Phen Green FL dye. Phen Green FL diacetate is a cell-permeable fluorescent dye quenched in the presence of Cu ions, and has been used effectively to image intracellular Cu ions in biological systems.^41^ Cells were incubated with Phen Green FL and imaged in a confocal microscope. Compared to the control (untreated cells), the Cu^2+^-treated cells showed a significant decrease in fluorescence intensity. When the Cu^2+^-treated cells were pre-incubated with melatonin, Phen Green fluorescence intensities afforded significant recovery and were similar to the Cu untreated controls **(Fig. 4G,4H,4I).** The recovery of Phen Green fluorescence upon melatonin treatment in Cu overloaded cells, along with the distinct Cu^2+^ selectivity of our chelators, indicates that these chelators will be able to chelate Cu from cells under oxidative stress conditions efficiently. Interestingly, we did not observe any significant difference in copper levels between cells treated with copper and cells exposed to copper preceding melatonin treatment in both WT HepG2 and ATP7B^-/-^ HepG2 **(Fig. 4J,4K).** indicating that though melatonin chelates copper and alleviates oxidative stress, it does not possibly facilitate copper export at the cellular level.

**Figure 4.**
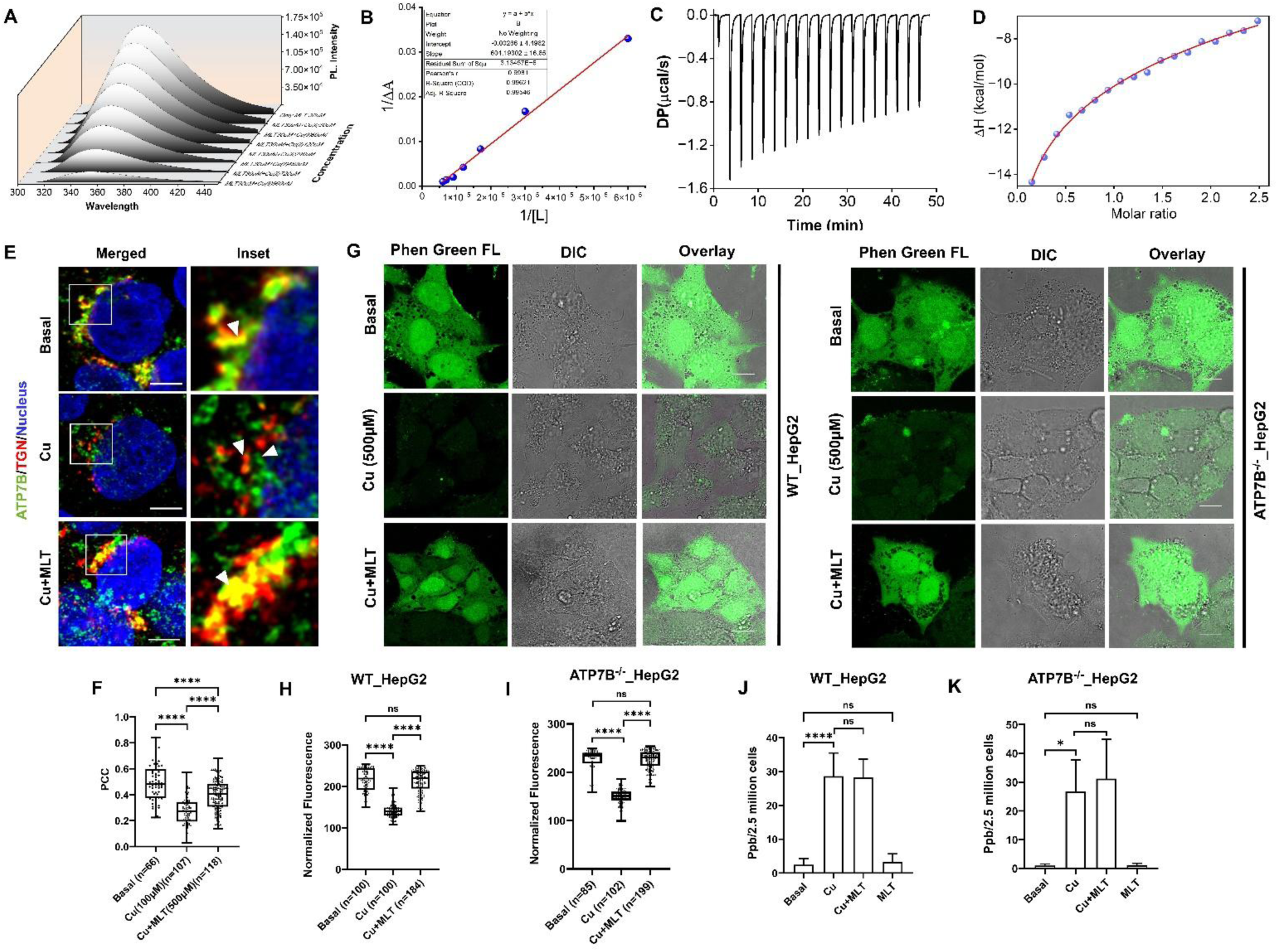
Melatonin binds copper selectively (in vitro and in vivo) and relocalizes ATP7B by chelating intracellular Cu^2^⁺. (A) Fluorescence emission spectra of melatonin (30 μM) upon incremental addition of Cu^2^⁺, showing dose-dependent quenching indicative of complex formation. (B) Benesi–Hildebrand plot assuming 1:1 stoichiometry yielded a binding constant (Kₐ) of ∼1.65 × 10^3^ M⁻^1^. (C) Representative ITC thermogram of melatonin titrated with Cu^2^⁺. (D) The binding isotherm derived from ITC data shows Kₐ ≈ 4.54 × 10^3^ M⁻^1^ with a stoichiometry of ∼1.59 and thermodynamic parameters ΔG ≈ –4.99 kcal/mol and TΔS ≈ –26.6 kcal/mol. (E) Immunofluorescence microscopy of ATP7B in WT HepG2 cells: in copper-treated cells, ATP7B vesicularizes; co-treatment with melatonin restores ATP7B localization to the trans-Golgi network (TGN). (F) Quantification of vesicular ATP7B redistribution and Phen Green FL signal intensities (mean ± S3; n ≥ 3). (G), (H), and (I) Phen Green FL signal is quenched in copper-treated cells and recovers upon melatonin pre-treatment. (J, K) ICP-MS analysis of total intracellular copper levels in WT and ATP7B⁻/⁻ HepG2 cells shows no significant difference between copper-only and copper + melatonin treated conditions (ns, p > 0.05).

Our findings establish melatonin as a selective and cell-permeable copper chelator that forms a stable complex with Cu^2^⁺ through amide nitrogen coordination. This dual role of melatonin—as both antioxidant and metal chelator—not only underscores its therapeutic promise in copper-overload disorders but also opens avenues for its rational repurposing in redox-linked pathologies.

### Melatonin Restores Redox Balance by Modulating Nrf2–HO-1 Signalling and Glutathione Oxidation

Based on our demonstration that melatonin directly scavenges reactive oxygen species (ROS) and chelates copper, we next sought to elucidate the molecular mechanisms by which melatonin alleviates oxidative stress in the context of Wilson disease (WD). Given that the nuclear factor erythroid 2-related factor 2 (Nrf2) pathway orchestrates antioxidant defence responses, we investigated melatonin’s role in modulating Nrf2 activation and its downstream effector, heme oxygenase-1 (HO-1). Under basal conditions, Nrf2 is primarily localized in the cytosol. However, under oxidative stress response, Nrf2 translocates to the nucleus, where it activates the transcription of antioxidant genes^42^. Using immunofluorescence assays, we observed that copper treatment markedly increased nuclear localization of Nrf2 in wild-type (wt) HepG2 cells, from approximately 12.5% under basal conditions to 59.65% following copper exposure **(Fig. S5A and S5B)**, indicating a robust oxidative stress response. However, pretreatment with melatonin reduced nuclear Nrf2 levels to 30%, suggesting that melatonin attenuates oxidative stress and thereby diminishes the requirement for Nrf2-driven antioxidant signaling. A similar pattern was seen in ATP7B^⁻/⁻^ HepG2 cells, where under basal conditions, nuclear Nrf2 levels were slightly elevated (∼19.5%) due to intracellular copper accumulation. Copper treatment further increased nuclear localization to ∼56%, which was significantly reduced to 25% upon melatonin pretreatment **(Fig. 5A** and **5B).** These findings reinforce the conclusion that melatonin’s antioxidant action effectively curtails oxidative stress, limiting the need for Nrf2 nuclear translocation^43^. Next, we checked the expression of Heme oxygenase-1 (HO-1). HO-1 is a cellular stress protein under the transcriptional control of Nrf2. In situations of elevated oxidative stress, haemoproteins can break down, releasing their heme component, which becomes "free heme" and enters the labile heme pool; this free heme is highly reactive and can directly trigger the production of reactive oxygen species (ROS), further amplifying the oxidative stress within the cell, potentially leading to cellular damage^44^. This leads to the expression of HO-1. HO-1 expression in wild-type and ATP7B^-/-^ cells was assessed by immunoblot analysis **(Fig. 5C** and **5D)**. Copper exposure significantly elevated HO-1 expression, consistent with Nrf2 pathway activation. Melatonin treatment reduced HO-1 levels, likely due to its direct antioxidant effects, which decrease the cellular demand for HO-1 upregulation. This indicates that melatonin enhances cellular antioxidant capacity and tunes antioxidant protein expression to maintain redox homeostasis.

**Figure 5.**
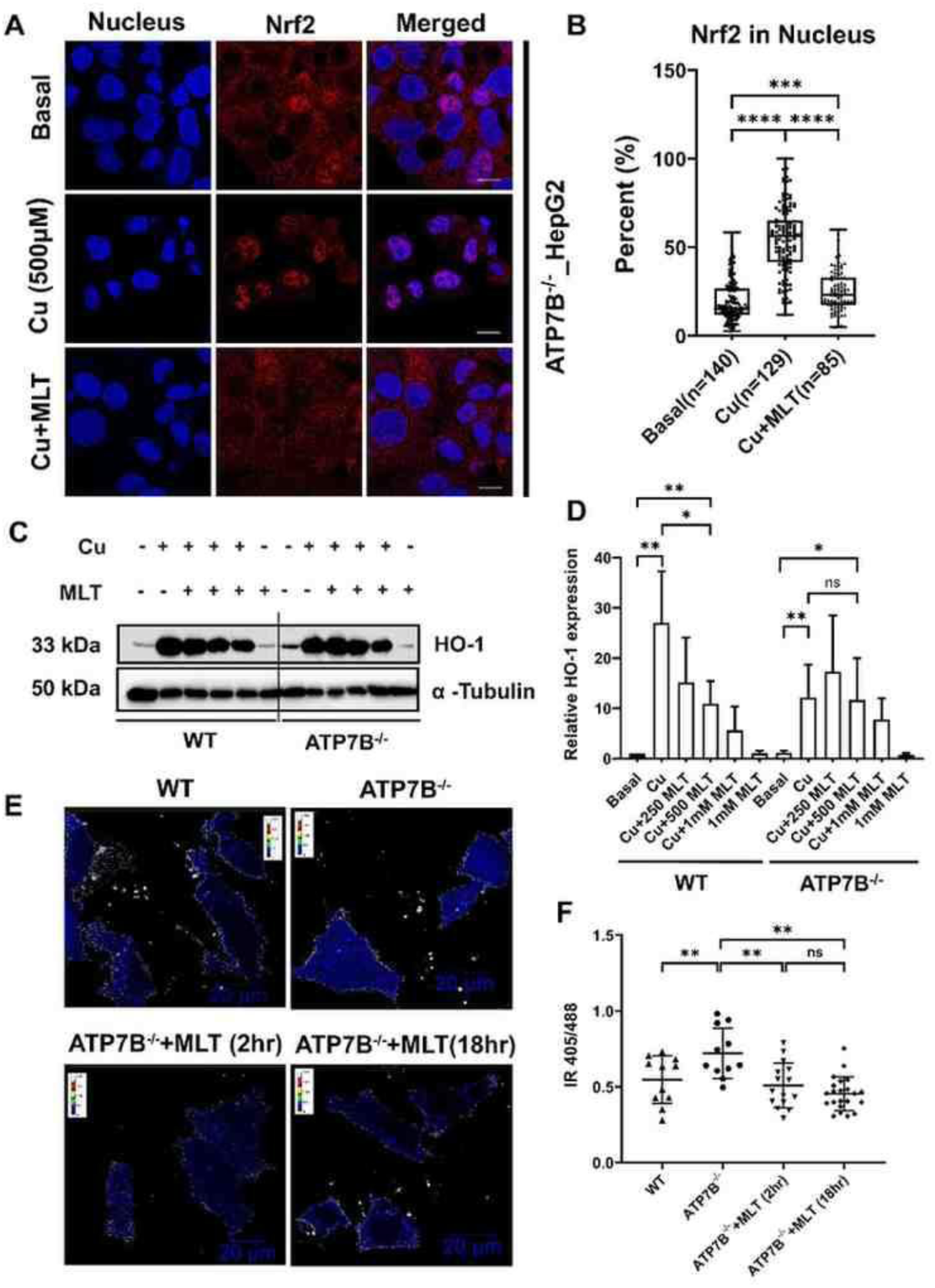
Melatonin Restores Redox Balance by Modulating Nrf2–HO-1 Signalling and Glutathione Oxidation. (A) Representative immunofluorescence images showing Nrf2 localization in ATP7B⁻/⁻ HepG2 cells under basal, copper-treated, and melatonin + copper conditions. Nuclei were counterstained with DAPI (blue); Nrf2 is shown in Red. (B) Quantification of nuclear Nrf2 localization in ATP7B⁻/⁻ in HepG2 cells. HepG2 cells reveal increased nuclear Nrf2 translocation upon copper exposure (∼56% in KO), which was significantly reduced by melatonin pretreatment (∼25% in KO) (mean ± SD, n=3, ***p < 0.001). (C) Western blot showing HO-1 expression in WT and ATP7B⁻/⁻ HepG2 cells under the indicated treatments. β-actin was used as a loading control. (D) Densitometric quantification of HO-1 band intensity normalized to β-actin. Copper exposure induces HO-1 expression, which is markedly reduced upon melatonin treatment (mean ± SD, n = 3, **p < 0.01). These results demonstrate that melatonin dampens copper-induced oxidative stress by inhibiting nuclear translocation of Nrf2 and downregulating its downstream target HO-1, thereby restoring redox homeostasis in both WT and ATP7B-deficient hepatic cells. (E) Pseudocolor ratiometric images of GRX1-roGFP2 are shown for WT and ATP7B ^(-/-)^ HepG2 cells (top panel), along with ATP7B ^(-/-)^ cells treated with melatonin for 2 hours and 18 hours, respectively (bottom panel). The images were acquired under live-cell conditions using a 63X oil immersion lens. Scale bar: 20 μM. (F) Quantitative analysis of the 405/488 ratio for GRX1-roGFP2 shows the following mean values: WT (0.5469, n=11), ATP7B ^(-/-)^ (0.7208, n=11), ATP7B(-/-) + MLT for 2 hours (0.5093, n=15), and ATP7B ^(-/-)^ + MLT for 18 hours (0.4541, n=22). Data are expressed as mean ± SD. Statistical significance: *****p < 0.0001, **p < 0.0015, *p < 0.0212, ns = 0.6767. Multiple comparisons were carried out using ordinary one-way ANOVA.

To further investigate whether deletion of ATP7B and subsequent melatonin treatment can decrease the GSSG/GSH ratio and alleviate oxidative stress. We performed ratiometric imaging of GRX1-roGFP2 under transient expression in the HEPG2 cells lacking ATP7B and treated with melatonin for 2 hours and 18 hours. We performed the ratiometric analysis of the GRX1-roGFP2 as an indicator of the GSSG/GSH ratio as described earlier.^45,46^ WT and ATP7B^-/-^ HepG2 cells were transiently transfected with GRX1-roGFP2 plasmids, and the Intensity Ratio of 405/488 (IR 405/488) was measured as described in the methods section **(Video1-4)**. WT cells displayed a mean IR405/488 of 0.5469, whereas ATP7B ^(-,-)^cells showed a significantly elevated ratio of 0.7208, consistent with increased glutathione oxidation (**Fig. 5E** and **5F**) due to copper accumulation. Notably, melatonin treatment for 2 and 18 hours significantly reduces the mean IR to 0.5093 and 0.4541 respectively, indicating that melatonin decreases GSSG/GSH ratio and effectively mitigates excess copper-induced glutathione oxidation in ATP7B^-/-^cells.

These results demonstrate that melatonin mitigates oxidative stress by modulating the Nrf2–HO-1 axis. Its ability to limit Nrf2 nuclear translocation and attenuate HO-1 expression under copper-induced stress highlights a regulatory role in redox homeostasis by scavenging free radicals, further reinforcing its therapeutic relevance in Wilson disease and other oxidative stress-related phenotypes.

### Melatonin mitigates copper-induced apoptotic cell death and improves mitochondrial health

Besides performing essential tasks, mitochondria are also a major site for the production of oxygen-based toxic species leading to apoptosis, the majority of which must be detoxified before they irreparably damage organelles and severely compromise ATP production.^47,48^ Previous studies have demonstrated that copper (Cu) exposure induces apoptosis, leading to significant cell death.^49^ To assess the protective effect of melatonin, we analysed the apoptotic cell population following Cu treatment in both wild-type (wt) and ATP7B^-/-^ HepG2 cells. In WT HepG2 cells, Cu treatment resulted in substantial cell death (22.8%), comprising 3.4% necrotic cells, 14.1% apoptotic cells, and 5.3% late apoptotic cells. However, melatonin pretreatment significantly reduces total cell death to 12.2%, with a decrease in necrotic (0.7%) and apoptotic (6.2%) populations, while the late apoptotic population remained unchanged (5.3%) **(Fig. 6A).** To evaluate melatonin’s protective role in a cellular model of WD, we examined its effect in ATP7B^-/-^ HepG2 cells, which exhibited heightened susceptibility to Cu-induced cytotoxicity. Following copper treatment, total cell death in ATP7B^-/-^ cells increased to 49.3%, consisting of 4% necrotic cells, 38.8% apoptotic cells, and 6.5% late apoptotic cells. Notably, melatonin pre-treatment significantly reduced cell death to 17.8%, with a decrease in necrotic (1.9%) and apoptotic (9.4%) populations, while the late apoptotic population remained at 6.5% **(Fig. 6B).** Total cell death in wt and HepG2 ATP7B^-/-^ cells has been analysed and represented in **Fig. 6C** and **Fig. 6D**. To probe the underlying mechanism of melatonin’s anti-apoptotic effect, we assessed mitochondrial oxidative stress using MitoSOX Red. This mitochondria-targeted fluorescent probe selectively detects superoxide (O₂•⁻), a primary ROS generated during mitochondrial dysfunction^50^. MitoSOX exhibits bright red fluorescence upon oxidation by superoxide, which we have quantified using flow cytometry. Following Cu treatment, we observed a three-fold increase in MitoSOX fluorescence intensity, indicative of elevated mitochondrial superoxide levels in both wt HepG2 and ATP7B^-/-^ HepG2 cells. Melatonin pre-treatment significantly reduced MitoSOX signal **(Fig.6E, 6F, 6G, and 6H)**, suggesting that melatonin effectively scavenges mitochondrial ROS. This decrease in mitochondrial oxidative stress correlates with the observed reduction in apoptosis, supporting a model where melatonin preserves mitochondrial integrity under copper overload. Together, these findings demonstrate that melatonin protects both WT and ATP7B^-/-^ HepG2 cells from copper-induced apoptosis by mitigating mitochondrial superoxide accumulation and preserving mitochondrial health.

**Figure 6.**
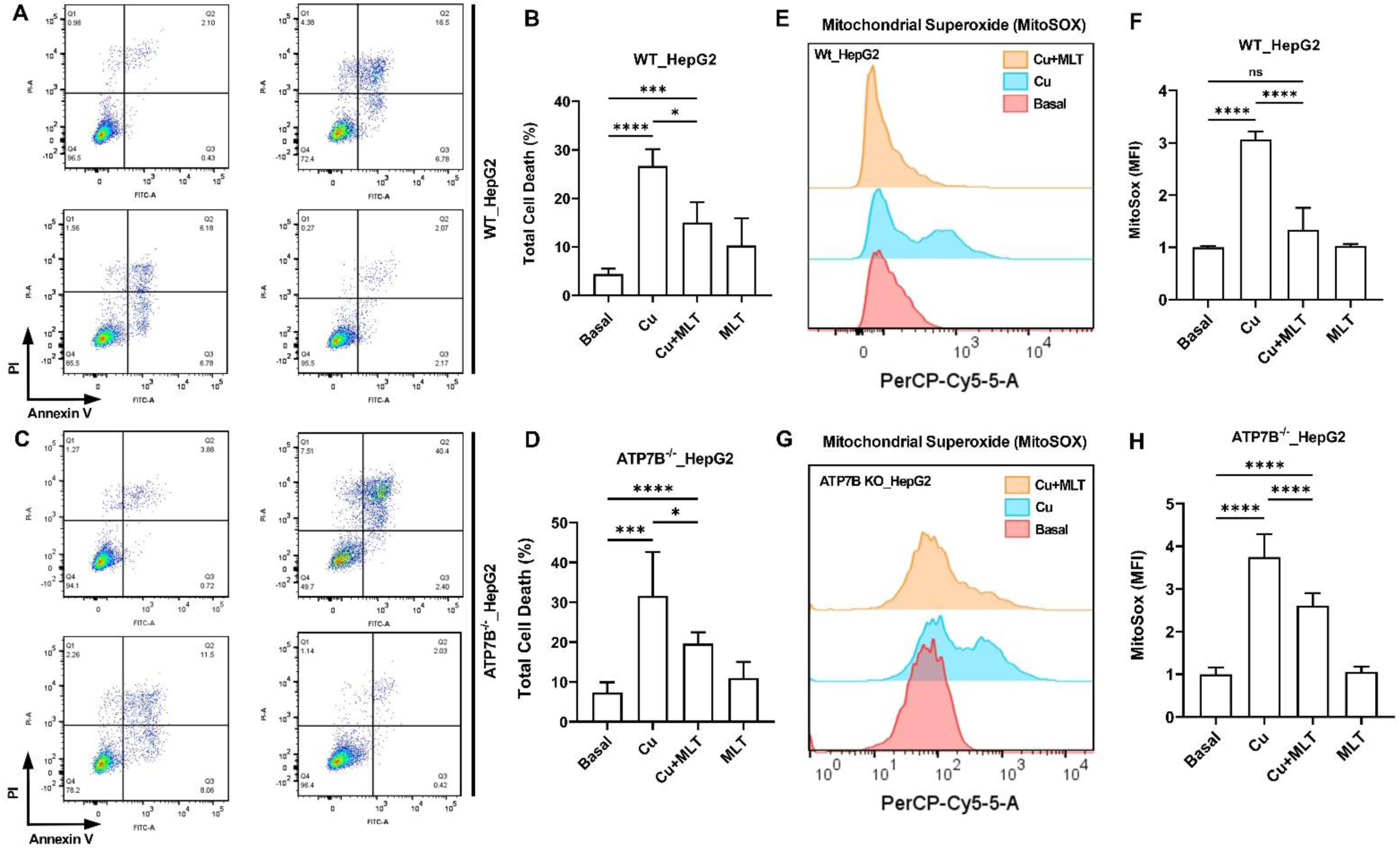
Melatonin protects WT and ATP7B⁻/⁻ HepG2 cells from Cu-induced apoptosis by reducing mitochondrial oxidative stress. (A) and (C) Flow cytometry analysis of total cell death in WT (A) and ATP7B⁻/⁻ (C) HepG2 cells using Annexin V–FITC and propidium iodide (PI) staining. Cu treatment increases total cell death in both cell types, with a marked rise in apoptotic populations. Melatonin pretreatment significantly reduces apoptosis and necrosis, while the late apoptotic population remains unchanged. (B) and (D) Quantification of the different cell populations (viable, necrotic, apoptotic, and late apoptotic) from panels A and B (mean ± SD, n = 3, **p < 0.01, ***p < 0.001). (E) and (G) Representative histograms of MitoSOX Red fluorescence in WT (E) and ATP7B⁻/⁻ (G) HepG2 cells showing elevated mitochondrial superoxide production upon Cu exposure and reduction following melatonin pretreatment. (F) and (H) Quantification of MitoSOX Red mean fluorescence intensity in WT (F) and ATP7B⁻/⁻ (H) cells under the indicated conditions (mean ± SEM, n = 3, ***p < 0.001).

### Melatonin reverses oxidative stress in a vertebrate model of Wilson disease

Building on the encouraging results from the cellular model and to extend our findings beyond cell-based systems, we investigated the therapeutic potential of melatonin using *in vivo* system by assessing its efficacy in counteracting copper toxicity in zebrafish embryos, an established vertebrate model for metal toxicity and oxidative stress studies^51^. We exposed 3-day post-fertilization (dpf) zebrafish embryos to 200 µM copper for 45 minutes (Cu) to induce oxidative stress (30 minutes of melatonin pre-treatment, 200 µM). Following copper exposure, oxidative stress levels were assessed using CellROX Orange, a fluorogenic probe widely used for measuring reactive oxygen species (ROS) in live cells and organisms^52^. Copper exposure significantly increased oxidative stress. We recorded a twofold increase in oxidative stress following Cu treatment, which was effectively reduced to basal levels after melatonin administration **(Fig. 7A** and **7B)**. We used flow cytometry to measure oxidative stress to further validate our microscopy findings. We recorded a significant decrease (∼75%) of Cell-ROX signal in copper-loaded cells treated with melatonin isolated from zebrafish embryo as compared to control ones (copper-treated) **(Fig. 7C** and **7D)**. In addition to oxidative stress, we assessed the extent of cell death in the embryos using annexin V-PI staining. Copper-treated embryos exhibited approximately 16% total cell death, which was reduced to 11% following melatonin administration **(Fig. 7E)**, further supporting melatonin’s cytoprotective effect.

**Figure 7.**
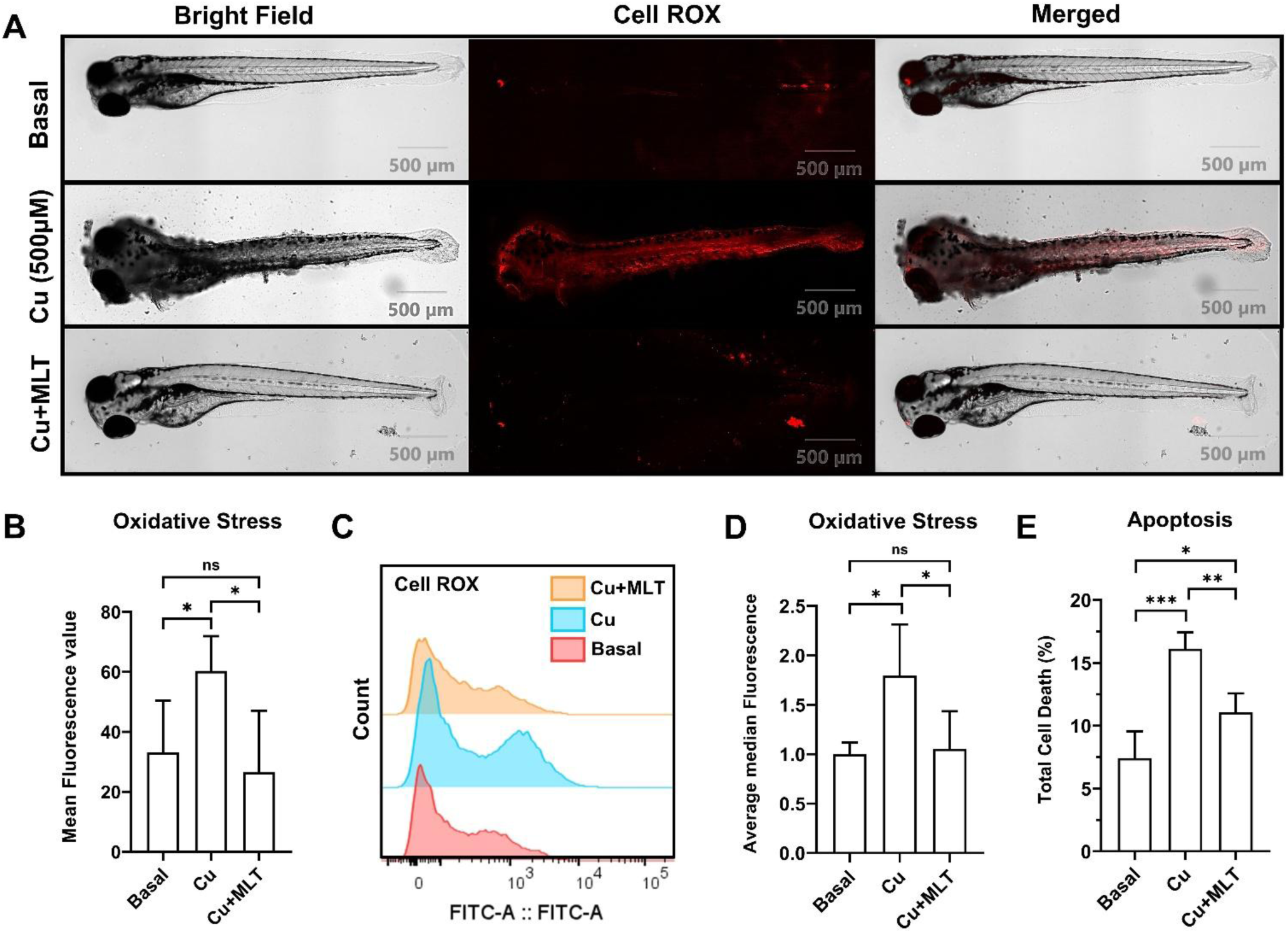
Melatonin mitigates copper-induced oxidative stress and cell death in vivo in zebrafish embryos. (A) Representative fluorescence images of 3 dpf zebrafish embryos stained with CellROX Orange to detect oxidative stress following treatment with vehicle (control), copper (200 µM, 45 min), or copper preceded by melatonin (200 µM, 30 min). Copper exposure markedly increases oxidative stress, which is suppressed to near-basal levels by melatonin pretreatment. (B) Quantification of CellROX fluorescence intensity in whole embryos (mean ± SD, n=6 embryos per group, ***p < 0.001). (C) Flow cytometry analysis of dissociated zebrafish embryo cells stained with CellROX confirms increased oxidative stress upon Cu treatment and its attenuation by melatonin. (D) Quantification of oxidative stress-positive cell populations from flow cytometry (mean ± SD, n = 3 independent experiments, ***p < 0.001). (E) Annexin V–FITC and propidium iodide (PI) staining of zebrafish embryos reveals that copper treatment induces ∼16% total cell death, which is reduced to ∼11% with melatonin pretreatment (mean ± SD, **p < 0.01).

These findings from the zebrafish model corroborate our cellular results, providing compelling evidence that melatonin mitigates copper-induced oxidative stress and preserves cellular viability in vitro and in a whole-organism context.

### Melatonin reduces oxidative stress and accumulated systemic copper in the *C. elegans* Wilson disease model

Following the assessment of the effect of melatonin in the zebrafish embryo model, we extended our investigation to determine whether melatonin could similarly alleviate copper toxicity in the *C. elegans* Wilson disease model (*cua-1^-/+^ C. elegans).* The N2 (wild-type) and tm12763 strains of C. elegans sourced from the National Bio-Resource Project (NBRP) were used for the study. Notably, the tm12763 worm carries a 63 bp deletion in exon 6 and a 53 bp deletion in the upstream intron of exon 6, resulting in the formation of a nonfunctional protein. *C. elegans* mutant strain (tm12763) lacking functional *cua-1*, the homolog of the human ATP7B, thereby serving as an adult model of Wilson disease. ^53^

Using the copper-sensitive probe CF4, we found that the tm12763 strain shows increased Cu-bound CF4 fluorescence intensity compared to the wild-type worms, localized in their intestinal region.^54^ **(Fig. 8A)** Quantitative analysis further revealed a significantly increased intensity of the Cu-bound CF4 in the *tm12763* worms compared to the wild-type worms. The increased intensity of Cu-bound CF4 was significantly attenuated with melatonin treatment in *tm12763* worms **(Fig. 8B).** The increased intensity of Cu-bound CF4 in *tm12763* worms reflects intracellular copper accumulation. To verify this, we performed ICP-MS analysis on whole adult worms. As anticipated, the *tm12763* worms exhibited an elevated amount of intracellular copper compared to wild-type worms. Melatonin treatment in *tm12763* worms decreases the amount of intracellular copper **(Fig. 8C).** To verify the specificity of CF-4 dye, we have also stained the wild type and cua-1 mutant strains with control CF-4 dye, which doesn’t give punctate-like staining like CF-4 and doesn’t respond to copper due to its modified isosteric receptor **(Fig. S6)**. Elevated intracellular Cu amount causes oxidative stress. To assess whether melatonin treatment can alleviate oxidative stress in *tm12763* worms caused by excess copper accumulation, we performed a CellROX assay. The results show that *tm12763* worms exhibit increased fluorescence signals consistent with high copper levels **(Fig. 8D).** Treatment with melatonin reduces the fluorescence intensity in these worms, indicating that melatonin can mitigate the oxidative damage caused by the excess accumulation of Cu **(Fig. 8E).**

**Figure 8.**
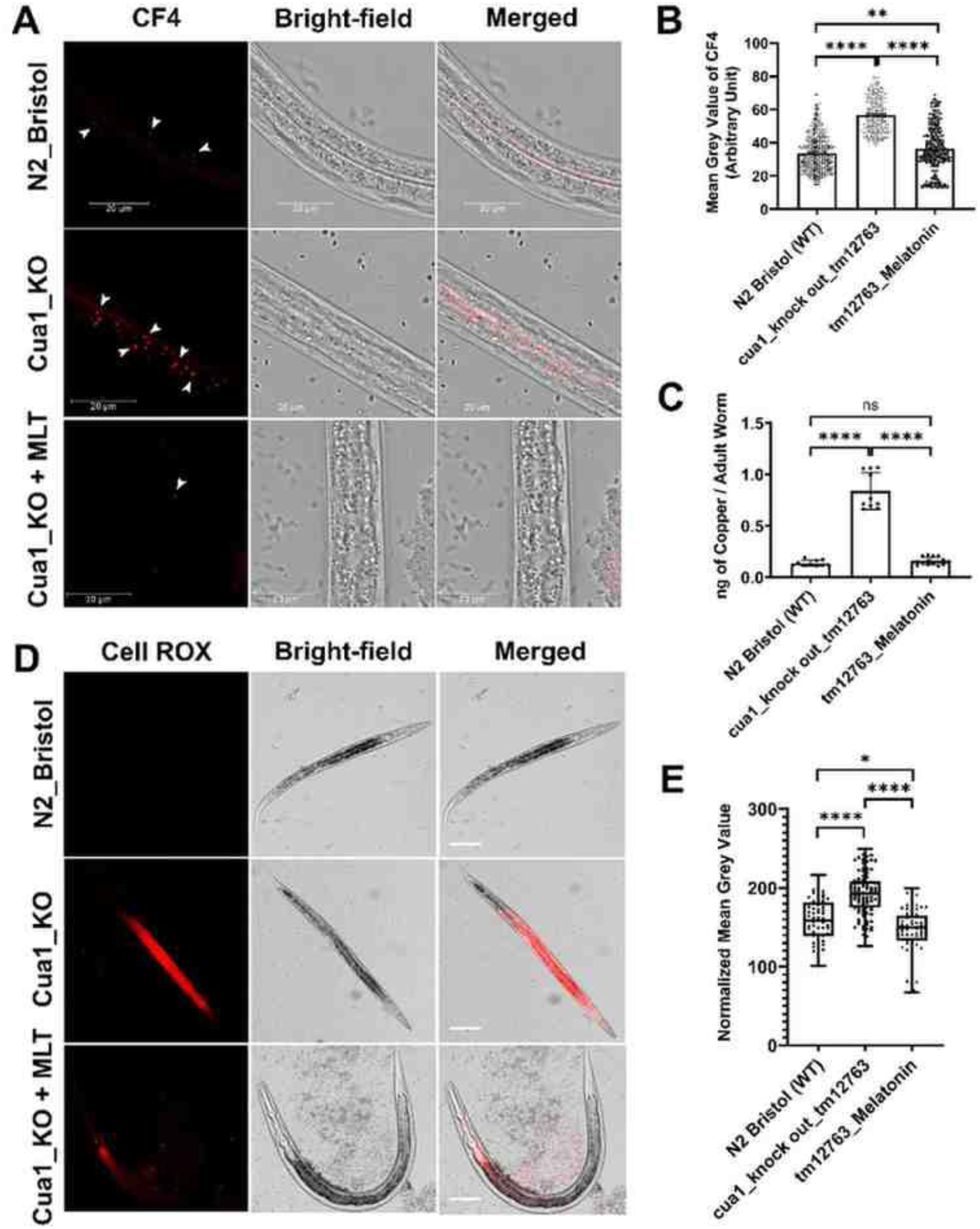
Melatonin treatment reduces copper accumulation and oxidative stress in the C. elegans cua-1 knockout model of Wilson disease. (a) Representative confocal images showing Cu-bound CF4 fluorescence localized to the intestinal region of L4-stage hermaphrodite C. elegans (N2 wild-type, tm12763 mutant, and melatonin-treated tm12763). Images were acquired using a 63× oil immersion objective. Scale bar: 20 μm. (b) Quantification of Cu-bound CF4 fluorescence intensity (Mean Grey Value, arbitrary units). tm12763 mutants exhibit significantly elevated fluorescence compared to wild-type, which is markedly reduced upon melatonin treatment (500 μM). Data represent mean ± SEM; n = 5 worms per group; one-way ANOVA; p = 0.0095, **p < 0.0001. (c) Intracellular copper content determined via ICP-MS in adult wild-type, tm12763, and melatonin-treated tm12763 worms. tm12763 worms show ∼6.13-fold higher copper accumulation than wild-type. Melatonin treatment reduces copper to 0.16 ng/worm, corresponding to a 5.36-fold decrease. n = 3 biological replicates; Student’s t-test; **p < 0.0001, ns = 0.9416. (d) Representative confocal images of CellROX Orange staining in wild-type, tm12763, and melatonin-treated tm12763 worms. Elevated ROS levels observed in tm12763 worms are markedly diminished following melatonin treatment (500 μM), indicating mitigation of copper-induced oxidative stress. Scale bar: 20 μm. (e) Quantification of CellROX fluorescence intensity (Mean Grey Value, arbitrary units) shows significantly elevated oxidative stress in tm12763 worms compared to wild-type, with a significant reduction upon melatonin treatment. Data represent mean ± SEM; n = 5 worms per group; one-way ANOVA; p = 0.0084, *p = 0.0006, ns = 0.4161.

Together, these results validate melatonin’s dual-mode function as a copper chelator and antioxidant in an adult Wilson disease model. Melatonin not only reduces copper overload but also alleviates the associated oxidative stress *in vivo*, mirroring its effects in cellular and embryonic models. The consistent efficacy observed across multiple biological systems highlights melatonin’s translational potential as a promising therapeutic candidate for Wilson disease.

### Melatonin-enclosed polymeric nanocapsule exhibits higher efficiency in alleviating oxidative stress

Using cellular and animal models, we demonstrated that melatonin is a promising candidate for WD therapy. However, a major limitation of melatonin as a therapeutic molecule is its short half-life in the bloodstream (45–60 minutes). It gets metabolized to 6-hydroxyderivative by CYP1A2 and eventually excreted in the urine, primarily as 6-hydroxymelatonin sulphate.^55^ We argued that encapsulated melatonin can circumvent the problem and ensure controlled and sustained release once inside the body. The nanocapsules were designed with a glutathione (GSH)-sensitive moiety, making them stimuli-responsive **(Fig. 9A, 9B)^56^**. Since oxidative stress leads to an increase in intracellular GSH levels, the nanocapsules rupture in response, enabling slow and sustained release of melatonin **(Fig. 9C)**. MNCs were incubated with glutathione to test the redox-triggered release of melatonin. HEPES buffer (pH 7.4) for 10 hours and subjected to transmission electron microscopy (TEM). TEM images showed a complete loss of the spherical morphology of the NCs (**Fig. 9C**), whereas control samples without GSH maintained their original spherical morphology. These observations support the GSH sensitivity of the nanocapsules and the disulfide bond cleavage mechanism. Melatonin-loaded MNCs were incubated with fetal bovine serum for 12 hours and analyzed using reverse-phase high-performance liquid chromatography (RP-HPLC) to assess whether encapsulation within MNCs enhances its stability. RP-HPLC analysis revealed that melatonin retained within the MNCs exhibited nearly a multiple-fold increase in stability compared to free melatonin (**Fig. 9D**). To assess the efficacy of MNCs in alleviating ROS, we measured oxidative stress levels after copper (Cu) treatment. wild-type HepG2 and ATP7B^-/-^ HepG2 cells were pretreated with varying concentrations of MNCs before being exposed to Cu. Our CellROX assay results showed that oxidative stress returned to basal levels after treatment with 250 µM of melatonin-loaded nanocapsules. In contrast, a three-fold higher concentration (750 µM) of free melatonin was required to achieve the same effect. This indicates that MNCs are significantly more effective than free melatonin in reducing oxidative stress **(Fig. 9E)^57^**.

**Figure 9.**
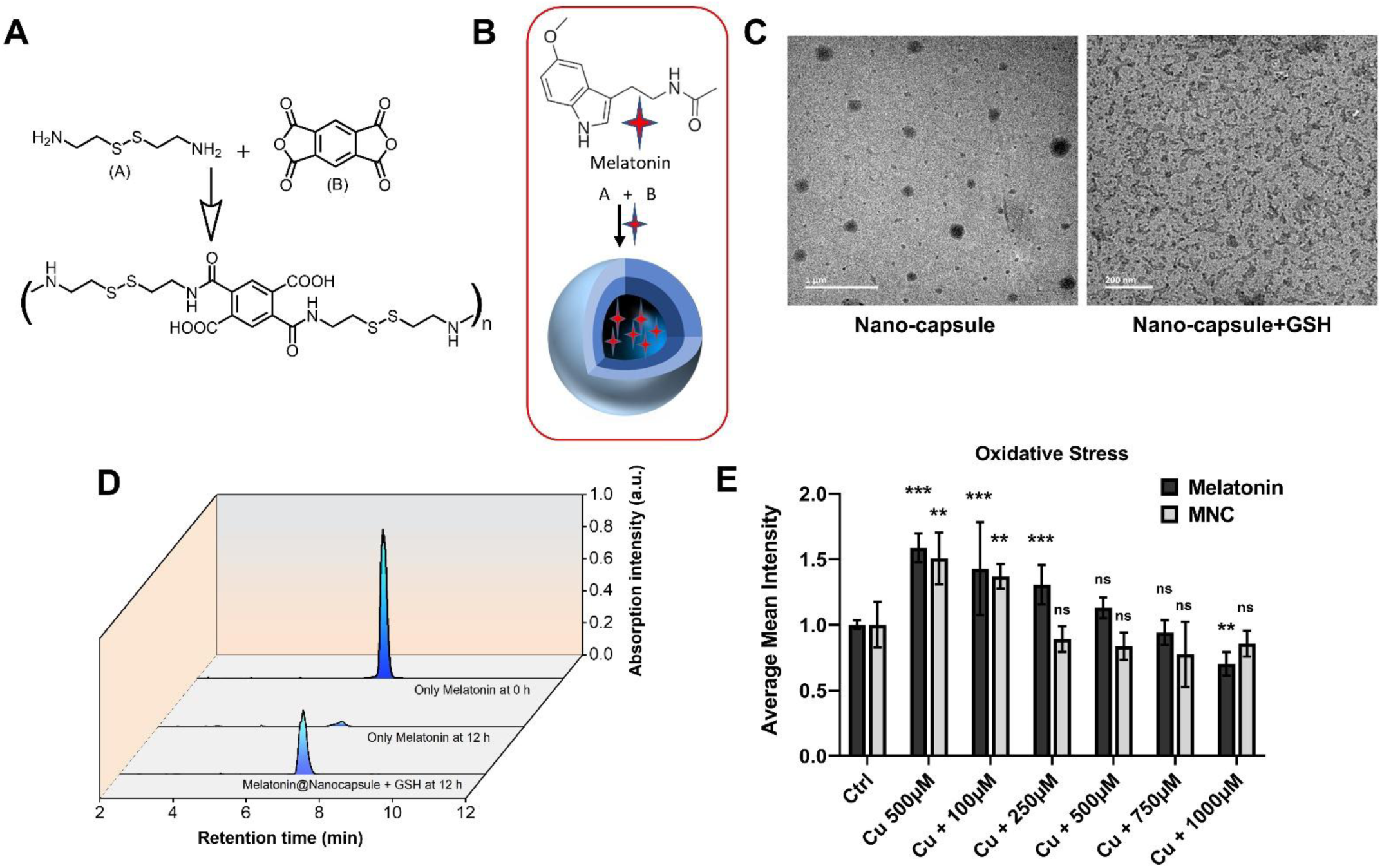
Melatonin-enclosed polymeric nanocapsules exhibits higher efficiency in alleviating oxidative stress. (a) Schematic illustration of GSH-responsive nanocapsules design for sustained intracellular melatonin release under oxidative stress. (b) Schematic of nanocapsules synthesis via interfacial inverse mini-emulsion technique involving amine–anhydride condensation. (C) Transmission electron microscopy (TEM) images of MNCs in the absence and presence of glutathione (GSH), demonstrating nanocapsules rupture upon GSH treatment, confirming glutathione sensitivity. (D). Reverse-phase HPLC analysis shows approximately ten-fold enhanced stability of encapsulated melatonin compared to free melatonin after 12 hours. (E) CellROX assay measuring intracellular oxidative stress in cells pretreated with MNCs or free melatonin prior to copper exposure. Treatment with 250 µM MNCs significantly reduced oxidative stress to basal levels compared to copper-only control, whereas free melatonin required a three-fold higher concentration (750 µM) to achieve similar effects. Data represent mean ± SD (n = 3). Statistical analysis was performed using an unpaired two-tailed t-test; p < 0.05, p < 0.01.

### Conclusion

In summary, we identify melatonin as a repurposed therapeutic that integrates Cu (II) coordination chemistry with intrinsic antioxidant activity to mitigate copper-induced oxidative pathology in Wilson disease models. Biophysical studies and molecular dynamics simulations revealed moderate Amide_N1_-Cu^2^ binding, complementing its radical-scavenging capacity and modulation of the Nrf2 pathway^44^. This dual mechanism restores redox balance while lowering copper burden in cellulo and in vivo. To address melatonin’s pharmacokinetic limitations^66^, we designed polyurethane–polythiourethane nanocapsules incorporating glutathione-cleavable linkers, enabling stimuli-responsive release under oxidative stress.^56^ Encapsulation extended melatonin’s half-life ten-fold and enhanced efficacy relative to the free molecule. Our findings have significant implications for the treatment of Wilson disease and other copper overload disorders such as Indian Childhood Cirrhosis (ICC), Idiopathic Copper Toxicosis, and Primary Biliary Cholangitis (PBC).^62^ These findings establish the first stimuli-responsive, dual-function therapeutic for copper overload disorders, showcasing how drug repurposing, molecular design, and advanced delivery platforms can converge to accelerate translational solutions.^57^

## Materials and Methods

### Cell lines and cell culture

HepG2 cells were grown and maintained in complete medium containing low glucose Minimum Essential Medium (MEM) (Himedia #AL047S) supplemented with 10% Fetal Bovine Serum (Gibco #10270-106), 1X Penicillin-Streptomycin (Gibco #15140-122).

### Zebrafish maintenance and handling

The Tübingen strain was used for all the experiments. All fish were maintained at 28.5 °C with a photoperiod of 14 h of light and 10 h of dark in a circulating water system. Zebrafish were fed dry food (Morning and evening) and live food (brine shrimp, Artemia sp.) (afternoon). Zebrafish were kept in specialised breeding tanks for embryo collection. In short, males and females were kept in breeding tanks separately by putting a separator in the tanks overnight. The separator was removed early in the morning, allowing them to interact and spawn for an hour. After an hour, the adults were removed and the embryos were collected using a mesh strainer, washed with and placed in a petri dish with E3 medium (5 mM NaCl, 0.17 mM KCl, 0.33 mM CaCl2, 0.33 mM MgSO4, 0.0001 % Methylene Blue) at 28.5℃ in an incubator. 3-day-old zebrafish embryos(3dpf) were exposed to the indicated CuCl2 solution for 45 minutes at 28.5 °C.

### C. elegans culture and strains

Wild-type animals used in this work were C. elegans variety Bristol, strain N2. The *cua-1* (tm12763) was obtained by the National Bioresource Project headed by the Mitani lab, Tokyo, Japan. The standard method of Brenner^65^ was used to culture synchronized *C. elegans* on Nematode Growth Media (NGM) plates at 20°C, which were inoculated with *E. coli* (OP50 strain) as a food source.

### MTT assay and Formazan quantification

3-(4,5-dimethylthiazol-2-yl)-2,5-diphenyl-2H-tetrazolium bromide (MTT) is used to evaluate the toxicity of Copper and Melatonin for all the experiments. Viability was calculated on the activity of NAD(P)H-dependent cellular oxidoreductases, which reduce the MTT dye into an insoluble formazan, which is purple and can be subsequently dissolved in dimethyl sulfoxide for colorimetric measurement. The cells were seeded into flat-bottomed 96-well culture plates (5000 cells/200 µl/well for the HepG2 cell line) and incubated in 5% CO_2_ incubator for 72 hours to allow the adherent cells to attach to the wells and let them grow up to 60-70% confluency. Treatment was given according to the experimental requirement and incubated for 24 hours. After gently pipetting off media, MTT (dissolved in PBS) was added to the MEM media (0.1 mg/ml), and the cells were incubated at 37°C, 5% CO2 for 4 hours. The medium was removed, and dark blue Formazan crystals formed by the cells were dissolved using 200 µl of DMSO. The absorbance was read at 570 nm wavelength on a multi-well plate reader. Data quantifications were done keeping relative viability with respect to the controls.

### Oxidative stress assay

CellROX Green was used to measure oxidative stress. CellROX Green is a fluorogenic probe, which is weakly fluorescent in its reduced state but exhibits bright green fluorescence upon oxidation by a free radical. The probe has an absorption/emission maxima of 508/525 nm, allowing precise quantification of intracellular reactive oxygen species (ROS) levels. HepG2 cells were cultured in 24-well plates and treated with the desired concentration of melatonin, followed by copper chloride to assess oxidative stress. After 24 hours of treatment, cells were incubated with 5*μM* CellROX Green reagent for ∼30 mins under standard culture conditions. The cells were then washed with 1X DPBS to remove excess dye, and the cells were detached by adding 100 μL 0.25% Trypsin-EDTA (Cat #25200072). After detachment, 300 μL of FACS buffer (1X DPBS, 2% FBS, 25 mM HEPES, 2 mM EDTA) was added to each well, and the samples were analyzed using flow cytometry BD LSR Fortessa (BD Biosciences) to quantify oxidative stress levels. Ten thousand cells were analysed for each condition. Mean intensity values were used for quantification. For live cell imaging (CellROX Green assay), cells were seeded on confocal dishes (Cat #200350 SPL Lifesciences). During imaging, cells were kept in imaging media containing phenol red-free low-glucose Minimum Essential Medium (MEM) (Gibco #41500-034) supplemented with 2% FBS, 20mM HEPES, and 1% Trolox (Sigma #238813). All images were acquired with the Leica SP8 confocal platform using an oil immersion 63X objective (NA 1.4) and deconvoluted using Leica Lightning software. Embryos were maintained in E3 medium to measure oxidative stress in Zebrafish embryos. After treatment completion, 20 *μM* CellROX orange was added and incubated for 30 minutes. Then, embryos were anaesthetized by adding Tricaine (Cat. # A-5040, Sigma) 0.003% in E3 Media (5 mM NaCl, 0.17 mM KCl, 0.33 mM CaCl2, 0.33 mM MgSO4, 0.0001 % Methylene Blue), and imaging was done using a spinning disk confocal with a 40 objective. A single cell suspension was made for flow cytometry, and the mean fluorescence intensity was analysed using a flow cytometer. ROS levels in *C. elegans* were measured under both normal conditions and following 24 hours of treatment with 500 μM melatonin. We used 10 μM Cell ROX orange (Invitrogen) to detect ROS levels *in vivo.* The OP50 around *C. elegans* was washed 3 times using M9 buffer; from this, 100 μL of precipitate was taken, and 100 μL of 5 μM cell ROX diluted in M9 buffer was added, and placed in a constant temperature at 20°C in the dark for 1 hour. *C. elegans* was then transferred to a 3% agar spacer with sodium azide and observed and photographed using a Leica confocal microscope.

### Staining of Labile Copper in *C. elegans*

Synchronized L4-stage worms were cultured on NGM plates seeded with *E. coli* OP50. After ∼3 days, *cua-1* knockout (*tm12763*) worms were treated with 500 μM melatonin (dissolved in M9 buffer: 22 μM KH₂PO₄, 42 μM Na₂HPO₄·2H₂O, and 85 μM NaCl) by adding the solution directly to the plates. Following 24 h of incubation, WT, *tm12763*, and melatonin-treated *tm12763* worms were washed twice with M9 buffer and incubated overnight with either CF4 or control CF4 dye. After staining, worms were gently washed with M9 buffer to remove excess dye and mounted onto 3% agarose pads on glass slides. For immobilization, 10 μL of 5 mM sodium azide was applied to the agarose pad. Coverslips were placed gently over the worms, and fluorescence images were acquired using a Leica SP8 confocal microscope with a 63× oil immersion objective and 1.28× digital zoom. Scale bar: 20 μm.

### Generation of ATP7B-/- HepG2

Guide RNAs(gRNA) targeted to ATP7B were cloned into pSpCas9(BB)-2A-Puro (PX459) using the Bbs1 restriction site. pSpCas9(BB)-2A-Puro (PX459) was a gift from Feng Zhang (Addgene plasmid #48139; http://n2t.net/addgene:48139) (Ran et al., 2013). Wild-type HepG2 cells were transfected with the clones using the polyplus jet-PRIME transfection reagent kit and kept until they reached 70% confluency. The concentration of puromycin was determined by keeping cells in a gradually increasing concentration in a 24-well plate. After 48 hrs of puromycin (3 μg/ml) treatment, the media was replaced with normal media to let the surviving cells grow to form colonies. Colonies were picked from Cloning cylinders (Merck C1059-1EA) and grown in a 24-well plate. The colonies were subsequently split into 12-well plates, 6-well plates, and 60 mm cell culture dishes. For characterisation, cells were stained for ATP7B in immunofluorescence. The sequence for gRNA for knocking out ATP7B is **CACCGCTATCGAGGCACTTCCACC** for sense, **AAACGGTGGAAGTGCCTCGATAGC** for anti-sense.

### Immunoblotting

After the respective CuCl_2_ and melatonin treatment, cells were pelleted down. Pellet was dissolved in 100µL of RIPA lysis buffer (10mM Tris-Cl, pH 8.0, 1.0% Triton X-100, 1mM EDTA, 0.5mM EGTA, 0.1% sodium deoxycholate, 0.1% SDS, 140mM NaCl, 1X protease inhibitor cocktail, 1mM phenylmethylsulfonyl fluoride) and kept for 30 minutes on ice. The solution is then sonicated with a probe sonicator (6 pulses, 5sec pulse on followed by 25 sec off,100mA). Bradford reagent (Cat #B6916, Merck) was used for protein quantitation, and 20 µg protein was loaded in each well. Protein sample was prepared by adding 4X NuPAGE loading buffer (Invitrogen#NP0007) to a final concentration of 1X and ran on 12% SDS PAGE. This was further followed by semi-dry transfer of proteins onto a nitrocellulose membrane (Millipore #IPVH00010). After transfer, the membrane was blocked with 3% skim milk in TBST (TBS with 0.1% Tween 20) for 3 hours at RT with mild shaking. Primary antibody incubation was done overnight at 4°C and then washed with 1X TBST (0.01% Tween-20). Secondary antibody incubation (Anti-rabbit HRP, Cat #7074 CST, 1:6000) was done for 1.5 hrs at RT, followed by TBST (3 times) and TBS (2 times) wash, and signal was developed by Clarity Max Western ECL Substrate (BioRad Cat #1705062) in ChemiDoc (BioRad). Details of Antibodies are as follows: α-tubulin (Affinity Biosciences Cat #AF7010, 1:20000), HO-1 (Affinity Biosciences Cat #AF5393, 1:1500), LC3B (Novus Cat# NB100-2220, 1:3000), Caspase 3 (Cat# ab90437).

### Immunofluorescence and microscopy

For fixed cell imaging, cells were fixed in 2% PFA in PBS solution for 20 min, followed by PFA quenching in 50mM NH4Cl solution for 20 min. Blocking was done in 3% BSA (bovine serum albumin) in PBSS (0.075% saponin in PBS) for 1.5 hours, followed by primary antibody incubation at room temperature for 1.5 hrs and in secondary antibody for 40 min. After subsequent washing with PBSS (3 times) followed by PBS (2 times), Coverslips were fixed on glass slides using SIGMA Fluoroshield™ with DAPI mountant. (Sigma #F6057). The solvent for antibody dilution was 1% BSA in PBS of ATP7B (Abcam #ab124973, 1:400), Golgin-97 (CST #13192, 1:400), and Nrf2 (Affinity Biosciences #AF0639, 1:200).

### Measurement of the GSH: GSSG ratio

Measurement of the GSH: GSSG ratio in wild-type HepG2 cells and in ATP7B^-/-^ HepG2, with or without melatonin treatment, was carried out as described earlier.^45^ Briefly, WT and ATP7B^-/-^ HepG2 cells were seeded on a glass-bottom dish (Cat #200350 SPL Lifesciences). After 24 hours, cells were transiently transfected with 1000 ng of GRX1-roGFP2 plasmid using polyplus jet-PRIME transfection reagent kit following the manufacturer’s protocol. The cells were then cultured overnight in MEM (Gibco) with or without 500 µM melatonin for 2 and 18 hours. Before imaging, the medium was replaced with DMEM without phenol red medium, again with or without 500 µM melatonin, and cells were incubated at 37 °C with 5% CO₂ for 2 hours. Sequential imaging was performed by exciting the sensor alternately at 405 nm and 488 nm, and emission was collected using a 500–554 nm band-pass filter in a Leica SP8 confocal microscope under live conditions. Generation of ratiometric images (IR405/488) and fluorescence intensity calculations at 405 nm and 488 nm has been performed in ImageJ Fiji software across 8 frames. The mean values were calculated for each cell and represented accordingly.

### Sample preparation for ICP-MS

HepG2 cells seeded on 60mm dishes were subjected to the appropriate treatment conditions at 70% confluency. Cells were then scraped with a cell scraper after being rapidly cleaned six times with 1X DPBS (Gibco #14200075). Cells were collected by centrifuging for five minutes at 2000 rpm. For counting of cells, the cell pellet was dissolved in 1ml of 1X DPBS and counted using Countess 3 Automated Cell Counters (Invitrogen). 2.5 million cells were taken and digested overnight at 95℃ in 100 µl of ICP-OES grade 65% HNO_3_ acid (Merck #1.00441.1000). 5 mL of 1X DPBS was added to each sample after digestion, and the samples were filtered through a 0.22-micron filter. Copper concentration was determined using an XSERIES 2 ICP-MS (Thermo Scientific).

For *C. elegans* copper measurement, a synchronized population of L4-stage worms was grown on OP50-seeded 35 mm NGM plates to the adult stage. 1 mL of 500 μM melatonin dissolved in M9 buffer was applied to the cua-1 strain 24 hours before harvesting. 30 synchronized young adult worms were then collected in 40 μL MilliQ ultrapure water. 100 μL of Nitric acid was added to the worms, and the samples were digested in a heating block set at 95°C overnight. The digested samples were diluted with 5 mL of MilliQ ultrapure water. Sample solutions were then thoroughly mixed and analysed. Values were normalized per young adult worm.

### Electron paramagnetic resonance (EPR) spectroscopy

Hydroxyl radical (•OH) generation was monitored via electron paramagnetic resonance (EPR) spectroscopy employing 5,5-dimethyl-1-pyrroline N-oxide (DMPO) as a spin-trapping agent. In the Cu^2^⁺/H₂O₂ system, a characteristic quartet EPR signal corresponding to the DMPO–OH adduct confirmed efficient •OH generation via Fenton-like redox cycling. Upon introduction of melatonin, a substantial attenuation of the DMPO–OH signal was observed, indicative of effective •OH suppression. This suggests that melatonin either directly scavenges •OH or modulates the Cu^2^⁺-mediated Fenton process by interfering with redox cycling and catalytic turnover.

### DPPH assay

The antioxidant property of melatonin was evaluated using the DPPH radical scavenging assay. DPPH is a stable free radical with an absorbance maximum at 517nm and undergoes reduction to DPPH-H in the presence of antioxidant molecules. DPPH absorbance was measured upon melatonin addition, keeping Ascorbic acid as a positive control, indicating its radical scavenging ability. The experiment was conducted in a 96-well plate format, with ascorbic acid as a positive control. The BioTek Cytation 5 Cell Imaging Multimode Reader instrument was used to measure the absorbance.

### 1H-NMR and DOSY NMR

For the ^1^H NMR titration, 8–10 mg of melatonin was dissolved in 600 μL DMSO-d₆. To this solution, 30 μL of a 1 M CuCl₂ solution in DMSO-d₆ was added. For Cu(I) studies, CuCl (as solid beads) was used as the Cu(I) source, and the entire procedure—including dissolution and mixing—was performed inside an argon-filled glove box to prevent oxidative degradation.

### Molecular Dynamics and DFT calculations

In the molecular dynamics system setup, initial structure of melatonin (CID 896) was obtained from PubChem and parameterized using Antechamber with GAFF2 atom types within the AMBER tools suite^67^. Initial configurations were prepared using PACKMOL^68^ with one melatonin and 64 Cu^2^⁺ ions. The system was solvated with TIP3P water and neutralized with 127 Cl⁻ ions in accordance with experimental stoichiometry. The AMBER simulation suite^69^ with the ff14SB force field was employed for the simulations. Energy minimization was performed using steepest descent, followed by switching to conjugate gradient after 500 steps for efficient convergence. The system was equilibrated for 0.5 ns at 300 K using Langevin thermostat. Pressure was maintained at 1bar using isotropic pressure coupling with a relaxation time of 1ps and a collision frequency of 2ps^-1^. Production MD was done for 500 ns under NPT ensemble. Electrostatics were treated using PME, and SHAKE was used for hydrogen constraints.

For DFT calculations, the level of theory employed to optimize all the geometries under study was M06-2X hybrid meta-exchange correlation functional^70^, along with 6-311++G(d,p) basis set. The PCM (Polarizable Continuum Model)^71-74^ implicit solvent model was used for the aqueous phase study. The minimum energy structures were confirmed by all positive values of consequent frequency calculations on the optimized geometries. The thermochemical results were obtained at 298.15 K and at 1 atm pressure. All electronic structure calculations were carried out using the Gaussian 16 software package^75^. Atomic population and second-order perturbation theory analysis were conducted using the NBO 3.1 module integrated within Gaussian 16.

### Fluorescence spectroscopy

All spectroscopic measurements were carried out in 20 mM HEPES buffer (pH ∼7.4) at room temperature using a Horiba Fluoromax spectrofluorometer. Melatonin was excited at 290 nm, and emission spectra were recorded over the 300–450 nm range. Metal ions (Cu^2^⁺, Ni^2^⁺, Ca^2^⁺, Mn^2^⁺, Zn^2^⁺, Mg^2^⁺, Na⁺, K⁺, and Cd^2^⁺) were introduced as their chloride salts. Upon gradual addition of each metal ion, no significant change in melatonin fluorescence intensity or spectral profile was observed, indicating a lack of strong interaction under these conditions.

### Mass spectrometry

Electrospray ionization mass spectrometry (ESI-MS) of melatonin (MW: 232.28 Da) incubated with CuCl₂ revealed prominent peaks at m/z 382.14, corresponding to melatonin–Cu complexes. These species are indicative of bis-adducts, suggesting that melatonin coordinates with copper ions, likely via multiple binding sites.

### Isothermal titration calorimetry (ITC)

ITC experiments were performed on a Malvern P analytical instrument at room temperature, using 19 successive injections at 180 s intervals. The syringe contained 1.0 mM CuCl_2_, and the sample cell contained a 75 µM aqueous melatonin solution.

### Assessment of cellular apoptotic status

To assess apoptotic status in HepG2 cells, cells were seeded in 12-well plates and pre-treated with 500 µM Melatonin for 3 hours, followed by 500 µM copper for 24 hours to induce apoptosis. After 24 hours, cells were detached using 0.25% Trypsin-EDTA to prepare a single-cell suspension, centrifuged at 600 rpm for 2 minutes, and the supernatant was discarded. Cells were washed twice with 1× phosphate-buffered saline (PBS) without calcium and magnesium, followed by resuspension in 1× Annexin V binding buffer. Annexin V Alexa Fluor 488 (Molecular Probes, A13201) was added as per the manufacturer’s protocol, and samples were incubated in the dark at room temperature for 30 minutes. Propidium iodide (PI; Sigma, Cat# P-4864-10ML) was diluted 1:10 in 1× Annexin V binding buffer, and 4 µL of the diluted PI solution was added to each sample, yielding a final concentration of 2 µg/mL. Samples were further incubated in the dark for 5 minutes at room temperature, washed with 500 µL 1× Annexin V binding buffer, and centrifuged again at 600 rpm for 2 minutes before final resuspension in 300 µL FACS buffer for flow cytometry analysis. In parallel, a single cell suspension was made from 72 hours post-fertilization (hpf) zebrafish embryos. This enables apoptosis assessment using a comparable Annexin V/PI staining protocol, ensuring consistency across the mammalian cell line and zebrafish models.

### Measurement of Mitochondrial Superoxide Radical

Mitochondrial superoxide was measured using MitoSOX Red. MitoSOX Red is a fluorescent probe that selectively targets and stains mitochondria. Upon oxidation by superoxide radical, MitoSOX Red exhibits fluorescence, allowing for both quantitative assessment of mitochondrial superoxide levels. After treatment completion (Melatonin and Copper), Cells were stained with 2.5 µM Mitosox Red (Cat#M36008, Invitrogen) reagent for 30 minutes. Cells were washed with 1X DPBS and kept in 100 µL Trypsin for 5 min. After that, cells were detached and suspended in 350 µL of FACS buffer (1XDPBS, 2% FBS, 25mM HEPES, 2mM EDTA) and were analyzed by flow cytometry BD LSRFortessa (BD Biosciences). 10,000 cells were analysed for each condition. Mean intensity values were used for quantification.

### Synthesis and characterization of Nanocapsules

Redox-responsive nanocapsules (MNCs) were synthesized via an interfacial inverse mini-emulsion technique by reacting cystamine and pyromellitic dianhydride in a stoichiometric ratio at the water-in-oil droplet interface, in the presence of melatonin as the encapsulated payload (**Fig. S7A**). The continuous phase was prepared by heating 10.0 g of cyclohexane and 120.0 mg of hypermer B246 at 50 °C until the hypermer B246 was fully dissolved. The dispersed phase contains 100 mg of cystamine, 5.0 mg of melatonin, and 1.0 g of 1 M KCl solution. The potassium chloride solution was employed to establish osmotic pressure within the droplets in the hydrophobic continuous phase, enabling the encapsulation of melatonin in the hydrophilic core.

The mixture containing the continuous and dispersed phases was left to pre-emulsify for 1 hour at 1000 rpm at room temperature. Afterwards, the mixture was sonicated using a Cole Parmer digital sonifier (USA) (1/4" tip) for 10 min (50s pulse; 50s pause) with an amplitude of 80 to achieve stable nanodroplets of the dispersed phase, while cooling the reaction mixture in an ice bath. An equimolar amount of pyromellitic dianhydride with respect to the cystamine was added to the additive phase containing 1.0 g of dichloromethane and triethylamine, serving as a base catalyst to facilitate the formation of nanocapsules. The latter was then slowly dropwise added to the emulsion and left to stir at room temperature for 4 hours to ensure complete polymerization. This results in the formation of nanocapsules with melatonin entrapped within the core. The reaction mixture was then filtered and redispersed in water. Redispersion was achieved by mixing 1g of the emulsion solution containing melatonin-loaded nanocapsules with 8 g of 1% SDS in water through effective stirring (∼1200 rpm) for 24 h at room temperature. Finally, the redispersed nanocapsule solution was passed through a 4μm filter paper to remove large aggregates and dialyzed for 24 h at room temperature. The morphology of the resulting nanocapsules were thoroughly characterized by dynamic light scattering (DLS), scanning electron microscopy (SEM), and transmission electron microscopy (TEM) **(Figure S7 B-E).** The effective encapsulation efficiency of melatonin in the nanocapsules was calculated by a previously reported protocol and found to be about 95% ^58,59^. Chemical analyses of the nanocapsules were performed using high-resolution solid-state ^13^C nuclear magnetic resonance (CPMS) and Fourier-transform infrared (FTIR) spectroscopy and X-ray photoelectron spectroscopy (XPS). SEM (**Fig. S7B, C**) and TEM (**Fig. S7D, E**) images indicated that nanocapsules had a spherical morphology with a mean diameter of 140 ± 10 nm with a narrower size distribution (Figure S6 B-E). The close examination of these nanocapsules under TEM (**Fig. S7D)** revealed a hollow interior with a thin shell (∼16.0 ± 4 nm thick), which agreed well with the results of previous reports. The DLS results suggested a mean size of 160 nm with a polydispersity index of ∼0.07, and this was consistent with TEM/SEM results. The surface potential was evaluated, and the zeta potential of −23.7 ± 3 mV was obtained. Further structural validation was provided by the FT-IR spectrum **(Fig. S7F)**, which displayed characteristic bands at 1633 cm⁻^1^ and 1521 cm⁻^1^, assigned to the stretching and bending vibrations of the C=O amide and N-H amide groups, respectively. Additionally, a band at 3520 cm⁻^1^ was observed, originating from the N-H amide vibration and the stretching vibration of the −OH group in the COOH moiety. The formation of the amide linkages through the amine-anhydride reaction was confirmed by analyzing the solid-state ^13^C NMR spectra of the nanocapsules. **Figure S7G** presents the NMR data obtained from the lyophilized nanocapsule sample. Aliphatic carbons were identified in the range of 25–49 ppm, while the ^13^C signals between 110–140 ppm correspond to the aromatic ring of pyromellitic dianhydride, which was used in the inverse-miniemulsion polymerization process. A distinct peak at 166 ppm was attributed to the C=O amide functionality, and a strong signal at 184 ppm confirmed the presence of a −COOH group. The XPS survey spectrum **(Fig. S7H-K)** revealed four prominent peaks corresponding to C 1s (284.9 eV), Pd 3d (340.61 eV), N 1s (399.11 eV), and O 1s (530.20 eV), confirming the presence of these elements in the HPOCs. The deconvoluted C 1s spectrum (Figure S6I) exhibited five distinct peaks: Csp^2^ (C=C, 283.5 eV), mixed carbons (C–C/C=N, 284.40 eV), Csp^3^ (C–N/C–OH, 285.50 eV), CCO (C=O, 287.35 eV), and Cπ→π* associated with π–π stacking interactions (289.16 eV). Elemental analysis of the nanocapsules, conducted using XPS, is illustrated in (Fig. S6H-K). The N 1s spectrum **(Fig. S7I)** displayed two peaks at 398.87 eV and 403.31 eV, corresponding to C–N linkages and protonated amine groups within the HPOCs. For O 1s, peaks at 529.88 eV and 532.36 eV were attributed to C=O and C–OH functionalities, respectively, while additional peaks at 531.23 eV and 534.58 eV were associated with hydrogen bonding and adsorbed water within the nanocapsules. The S 3d spectrum showed two peaks at 337.65 eV and 342.74 eV, corresponding to S (II) 3d₅/₂ and S (II) 3d₃/₂, respectively, indicating the preservation of disulfide linkages.

To evaluate the redox-responsive behaviour of the disulfide linkages within the nanocapsules, melatonin-loaded MNCs were incubated with glutathione (GSH) at a concentration of 1× PBS buffer (pH 7.4). The GSH-induced release of melatonin was monitored via the increase in its characteristic luminescence (λ_Abs_^Max^ = 305 nm; λ_Ems_^Max^ = 350 nm). Results showed that ∼95% of melatonin was released within 8 hours. In contrast, control experiments in the absence of GSH showed negligible release, confirming that the reductive cleavage of disulfide bonds by GSH is essential for melatonin release. The sustained release of melatonin over 8 hours also underscores its potential for controlled drug delivery, a critical factor in enhancing therapeutic efficacy **(Fig. S7, L).**

### Dynamic Light Scattering (DLS)

DLS studies were performed using a Brookhaven Instruments ZetaPALS with a 4.0 mW HeNe laser (λ = 633 nm) with a scattering angle of 173 at 25 °C. Nano-software was used to evaluate the size distribution following a non-negative least-squares (NNLS) analysis.

### Fourier Transform Infrared (FT-IR) Spectroscopy

The FT-IR spectroscopic measurements were carried out using a PerkinElmer G spectrophotometer. The spectra were recorded in the 400− 4000 cm−1 range in a KBr pellet.

### High-Resolution Solid-State Nuclear Magnetic Resonance (NMR) Spectroscopy

The High-Resolution Solid-State NMR measurements were performed in a Bruker Avance II (500 MHz) spectrometer (9.4 T wide-bore magnet). The signal for the aromatic carbon of hexamethylbenzene appeared at 132.1 ppm, which was used as the standard for calibrating the carbon chemical shift scale following the Hartmann−Hahn condition (1H = H B1H = C= 1C) for cross-polarization (CP).

### Electron Microscopy Imaging

A JEOL JEM 2100 microscope operated at 200 kV was used for recording TEM images. For studies, the nanocapsule solution was drop-casted on the TEM grids (lacey carbon Formvar-coated Cu grids (300 mesh)). No additional staining reagent was used for TEM analysis. Scanning electron micrographs (SEM) of secondary electrons (SE) were recorded using a ZEISS SUORA 35 VP scanning electron microscope.

### X-ray photoelectron spectroscopy (XPS)

Chemical composition of the nanocapsule was carried out by X-ray photoelectron spectroscopy (XPS) using the Omicron Nanotechnology spectrometer with 300 W monochromatic AlKα X-ray as an excitation source. The survey and core-level XPS spectra were recorded from at least three different spots on the samples. The analyzer was operated at a constant pass energy of 20 eV, with the Cls peak set at BE 285 eV to overcome any sample charging. For the XPS elemental analysis, freestanding nanocapsules were transferred onto a PLATYPUSTM gold-coated silicon wafer and dried at room temperature. CasaXps software was used for data processing.

### In Vitro Release Kinetics

The melatonin-loaded nanocapsules, prepared by the inverse-mini-emulsion technique, were redispersed in HEPES buffer, and the glutathione-mediated release of the encapsulating melatonin as a function of time was assessed using fluorescence spectroscopy (for melatonin λ_Max_Abs = 305 nm, λ_Max_Ems = 350 nm). The extent of melatonin released (in percentage) as a function of time was evaluated with respect to the amount of melatonin that was originally encapsulated in nanocapsules. All fluorescence studies were performed using a PTI-Horiba QuantaMaster 400 fluorometer. The fluorescence intensity of the reference sample having only free melatonin was set as 100% (F^0^). The quantity of melatonin released due to the rupture of the nanocapsules was assessed by calculating the difference in F^0^ and the fluorescent intensities of melatonin that was released from nanocapsules after centrifugation (10,000 rpm, 40 min, 0 °C). An average value of three identical experiments was used for all of our studies. The encapsulated amount was then calculated for each sample and taken as 100% for the release studies of the respective samples.

### TEM of Nano capsule- GSH

For TEM analysis, 1 mg of the nanocapsule was dispersed in 1 mL of Milli-Q water and drop-cast onto a copper TEM grid. Subsequently, 50 μL of a 10 mM glutathione (GSH) solution was added to the same solution, and the sample was incubated overnight at 37 °C to allow interaction. Post-incubation, the treated sample was again drop-cast onto a fresh copper grid for imaging.

### Melatonin Loading and Release Studies

Fluorescence of 100 μM melatonin in distilled water was recorded. Nano capsules were dispersed in the same solution, and fluorescence was measured at defined time intervals. A gradual decrease in fluorescence intensity indicated melatonin encapsulation, and the percentage loading was calculated after 12 h based on fluorescence quenching. For release studies, glutathione (GSH) was added to the melatonin-loaded nano capsule dispersion, and fluorescence recovery was monitored over time. A progressive increase in fluorescence intensity confirmed melatonin release, from which the release efficiency was quantified.

### HPLC Analysis of Melatonin Stability

1 mg of melatonin was dissolved in 1 mL of methanol and analyzed by reverse-phase HPLC (Waters system) using an acetonitrile–water mixture as the mobile phase. The chromatograms were recorded at 0 h and 12 h to monitor the stability of melatonin. Separately, melatonin-loaded nanocapsules were treated with glutathione (GSH), and stability was evaluated using the same HPLC gradient conditions. The analysis was performed on a reverse-phase C18 column, and melatonin elution was monitored at 278nm.

### Image analysis and statistics

Images were analysed in batches using ImageJ/Fiji, an image analysis software. For the colocalization study, the Colocalization Finder plugin was used. ROIs were drawn manually on the best z-stack for each cell. Pearson correlation coefficient (PCC) was used to quantify colocalization. For protein co-localization studies, to calculate the trans-Golgi network (TGN) distribution of ATP7B, fractions of ATP7B localized in Golgin-97 marked compartments were obtained using the Pearson correlation coefficient (PCC). Nrf2 localization inside the nucleus is quantified after splitting the cells into RGB channels and drawing ROIs manually around the nucleus region. For the box plots, the box represents the 25th–75th percentiles and the median in the middle. The whiskers show the data points within the range of 1.5 × interquartile range from the first and third quartiles. Non-parametric tests for unpaired datasets (non-parametric Mann–Whitney *U* test/Wilcoxon rank-sum test) were performed for all the samples; \**p* < 0.05; \*\**p* < 0.01; \*\*\*\**p* < 0.0001; ns, not significant. Cu-bound CF4 fluorescence was quantified as described in Chakraborty et al. (*Metallomics*, 2023).^68^ Confocal images were analyzed using ImageJ (Fiji). Images were scaled to micrometres and converted to 8-bit format, followed by auto-thresholding using the Otsu method. Intestinal regions were selected as ROIs, and the “Analyze Particles” tool was used to measure fluorescence intensity. GraphPad Prism was used for statistical analysis and plotting. ImageJ macro codes for image analysis are available at <u>GitHub</u>.

### Animal ethics statement

Zebrafish were bred and maintained, and experiments were performed in accordance with guidelines and protocols approved by the institutional animal ethics committee of the Indian Institute of Science Education and Research, Kolkata. All works performed with Zebrafish and *C.elegans* were in accordance with the Guidelines on the Handling and Training of Laboratory Animals by the Indian Institute of Science Education and Research, Kolkata.

### Safety

There are no unexpected, new, and/or significant hazards or risks associated with this work.

## Acknowledgment

This work was supported by DBT-Wellcome Trust India Alliance Fellowship (IA/I/16/1/502369) and Department of Science and Technology, Government of India, (DST/TDT/TC/RARE/2022/32), Govt. of India, IISER K intramural funding to A.G. and STARS-2 Grant (2023-0210) from Ministry of Education to A.G. and N.S. A.D. acknowledges the ANRF-J.C. Bose Fellowship and funding through the JBR/2023/000005 grant. NS acknowledges computational resources from PARAM Shakti and PARAM Rudra facilities established by CDAC, under the MeitY. C.J.C. was supported by NIH grant GM79465.

R.P. is supported by a pre-doctoral fellowship from the Department of Biotechnology, Government of India, India. and S.M. and R.B. were supported by a pre-doctoral fellowship from the Council of Scientific and Industrial Research, S.S., S.D., S.P., and D.B. were supported by a pre-doctoral fellowship from the University Grant Commission, Govt of India. RR acknowledges IISER Kolkata for providing a Postdoctoral fellowship. The authors also acknowledge the IISER Kolkata Central Instrumentation Facility (iCIF) for support and instrumental facilities.

We would also like to thank Tamal Ghosh for helping in flow cytometry experiments at the Department of Biological Science (IISER Kolkata) and Dr. Santosh Ch Das (Department of Earth Sciences, IISER Kolkata) for helping in using ICP-MS at the Department of Earth Science (IISER Kolkata). We thank Moulinath Acharya (the National Institute of Biomedical Genomics) and Santanu Saha Roy for helping us with the Zebrafish experiments.

## Author contributions

RR, ANR, SS, KC, RB, Pruthwiraj, SM, ATP and DB conducted the wet lab experiments and analyzed the data. SK helped with the data analysis RR, SP did the MD-simulation runs. CJC, AB, NS, AD and AG planned the experiments. NS, AD and AG wrote the manuscript. All authors reviewed the results and approved the final version of the manuscript.

## Conflict of interest

The authors declare that they have no conflicts of interest with the contents of this article.

## Notes

### Competing Interest Statement

The authors have declared no competing interest.

