## Supplementary material for "Repurposing Melatonin in dual-mode for Wilson disease therapy as a Copper Chelator and an antioxidant agent": Supplemental figures figure biorxiv.pdf

**Supplementary figures**

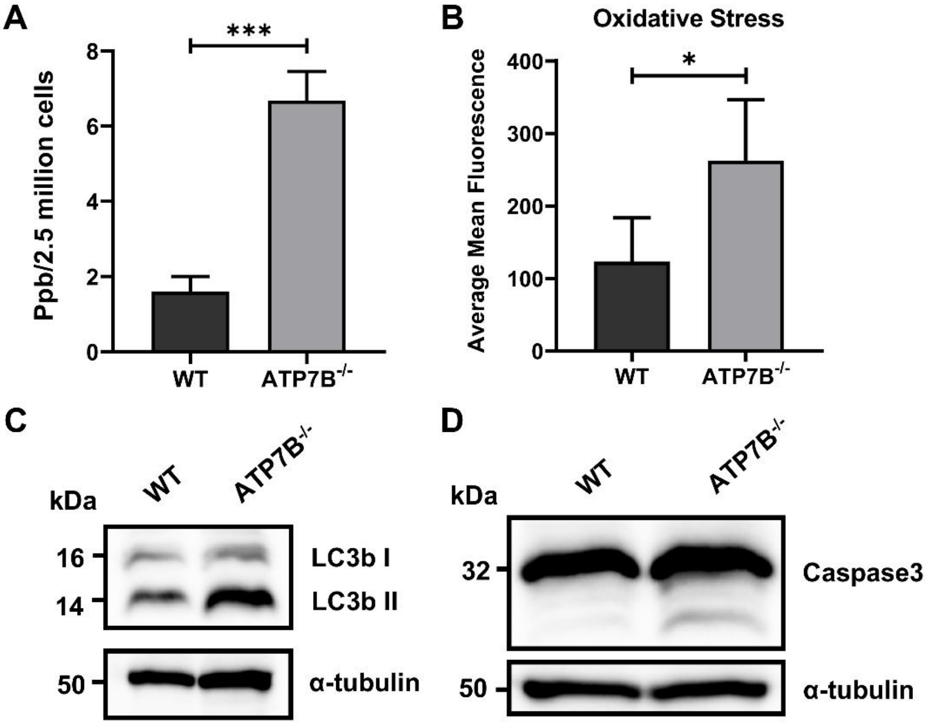


**Figure S1.**

**ATP7B deficiency leads to elevated intracellular copper, oxidative stress, and activation of apoptosis and autophagy pathways.**

(A) Quantification of intracellular copper levels in wild-type and ATP7B–/– HepG2 cells using Inductively Coupled Plasma Mass Spectrometry (ICP-MS). KO cells show ~4-fold higher copper accumulation compared to wild-type ($p < 0.01$, unpaired t-test, $n = 3$). (B) Intracellular ROS levels in ATP7B–/– and wild-type HepG2 cells under basal and copper-treated conditions (500 µM, 24 h), measured using CellROX green reagent. Copper-induced oxidative stress is significantly higher in KO cells ($p < 0.001$, $n = 3$). (C) Immunoblot analysis of LC3 lipidation, indicating conversion from LC3-I to LC3-II in ATP7B–/– cells, consistent with autophagy activation under copper stress. α-tubulin was used as a loading control for all immunoblot analyses. (D)Immunoblot showing cleavage of Caspase-3, a hallmark of apoptosis, in ATP7B–/– cells upon copper exposure.  Data represent mean ± SD from three independent experiments. Statistical analysis was performed using an unpaired t-test. *$p < 0.05$; **$p < 0.01$; **$p < 0.001$.

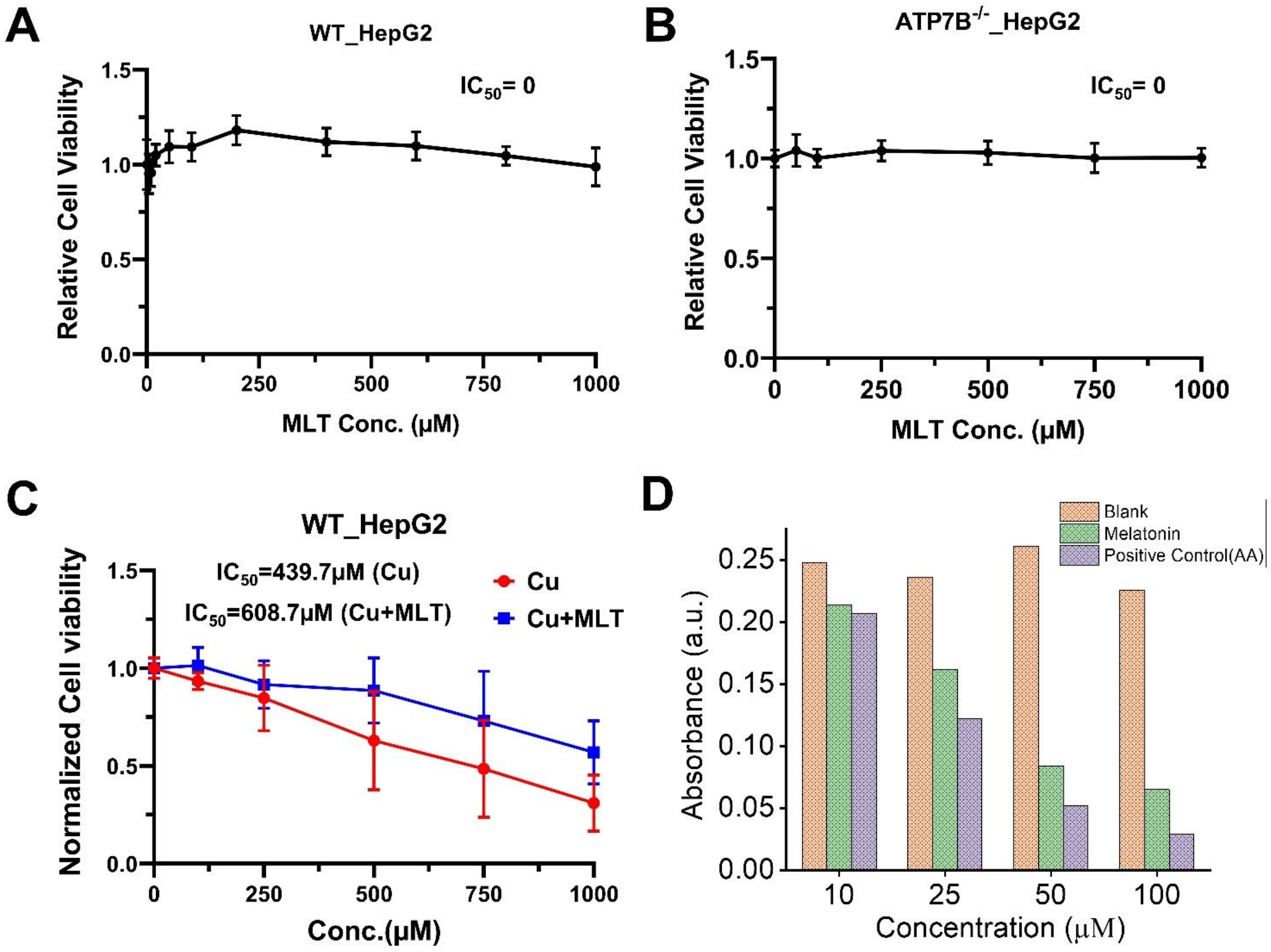


**Figure S2.**

**Melatonin is non-toxic and exhibits potent antioxidant activity contributing to its cytoprotective effect against copper toxicity.**

(A, B) Cell viability of wild-type and ATP7B–/– HepG2 cells after 24 h melatonin treatment (0–1 mM), assessed using MTT assay. Melatonin shows no cytotoxicity up to 1 mM ($n$ = 3). (C) Dose–response analysis of copper-induced cytotoxicity in WT HepG2 cells with or without melatonin pre-treatment. $IC_{50}$ increases from 964 μM to 14 mM after melatonin treatment, indicating a strong protective effect ($n$=3). (D) DPPH free radical scavenging assay comparing melatonin with ascorbic acid. Melatonin exhibits significant, dose-dependent antioxidant activity, confirming its intrinsic free radical scavenging potential. All data are presented as mean ± SD from three or more independent experiments.

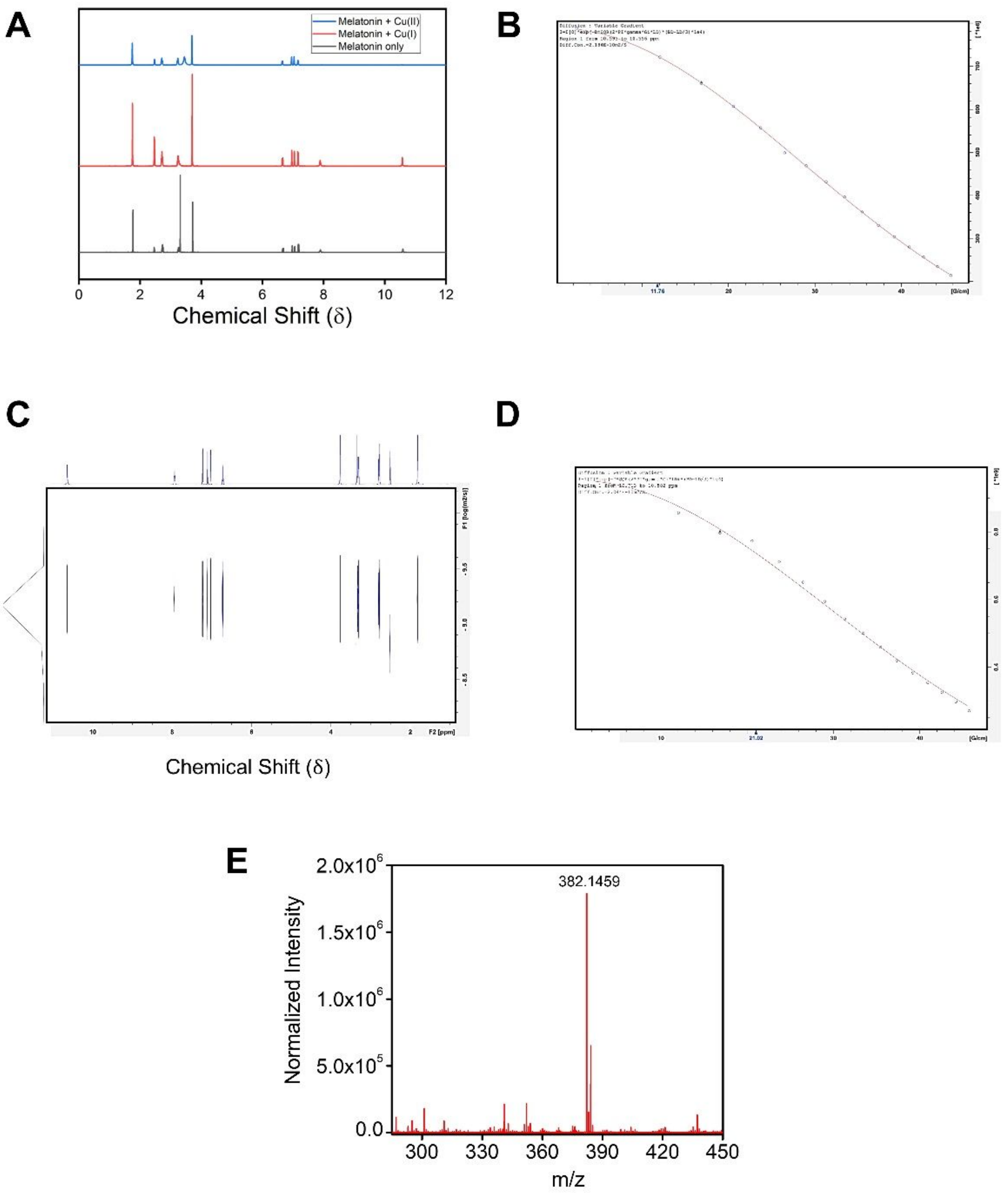


**Figure S3.**

**Spectroscopic evidence supporting melatonin–copper interaction.**

(A) Full 1H-NMR spectra of melatonin in the absence and presence of $Cu^{2+}$ and $Cu^{1+}$, highlighting key chemical shift changes. Compared to the representative spectra shown in the main Figure 3A and 3B (aromatic and aliphatic regions, respectively), this panel presents the complete spectral window. The disappearance of the amide proton and downfield shift of surrounding signals indicate specific coordination of $Cu^{2+}$ to the amide nitrogen of melatonin. (B) DOSY (Diffusion-Ordered Spectroscopy) spectra of melatonin-copper complex. (C) DOSY (Diffusion-Ordered Spectroscopy) plot of free melatonin. (D) DOSY spectra of the free melatonin, showing altered diffusion behaviour. The diffusion coefficient increases from $2.044 \times 10^{-10}$ $m^2/s$ (melatonin) to $2.184 \times 10^{-10}$ $m^2/s$ upon complexation, consistent with a change in hydrodynamic size due to copper binding. (E) Mass spectrometric analysis of the melatonin–copper complex. The appearance of a peak at m/z 382 corresponds to a complex comprising two copper ions coordinated with melatonin, suggesting a 2:1 stoichiometry.

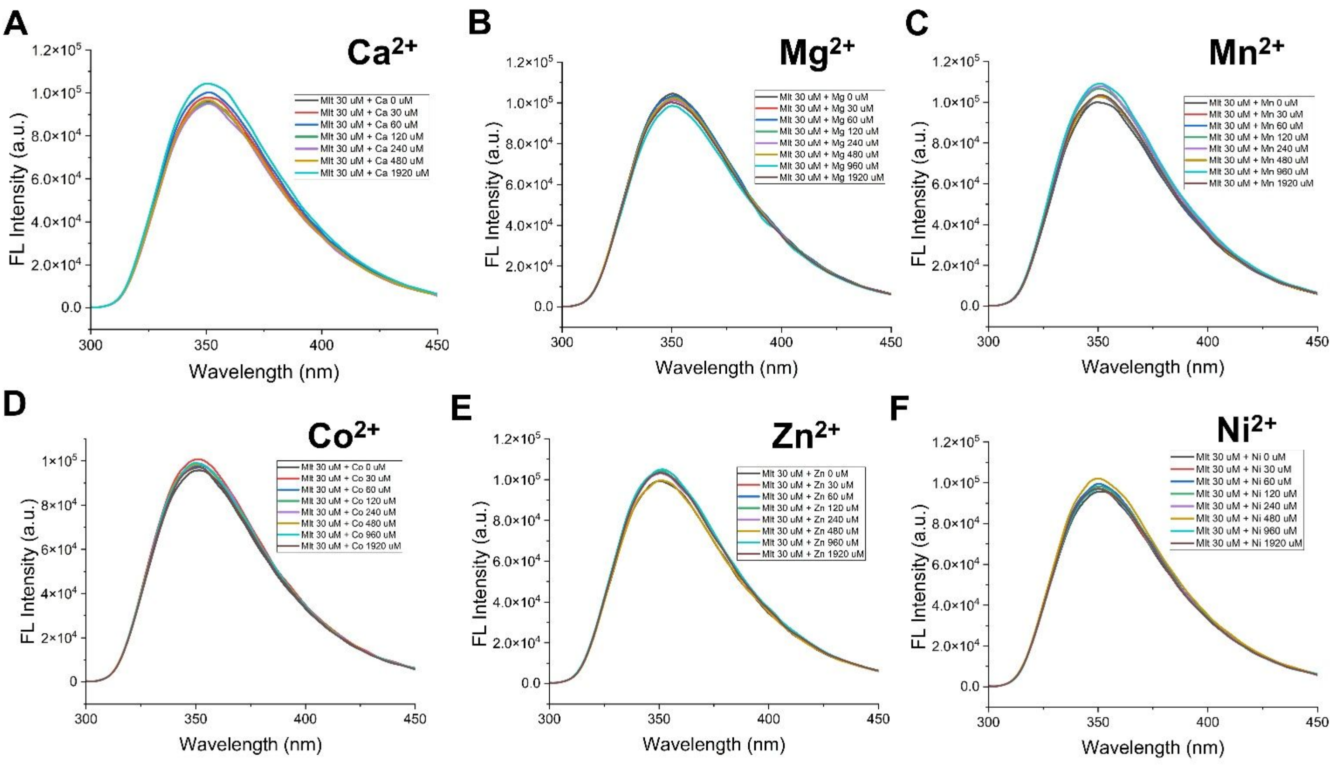


**Figure S4.**

**Melatonin selectively binds $Cu^{2+}$ but not other divalent metal ions**
(A–F) Fluorescence emission spectra of melatonin (30 µM) upon titration with various divalent metal ions—$Ca^{2+}$ (A), $Mg^{2+}$ (B), $Mn^{2+}$ (C), $Co^{2+}$ (D), $Zn^{2+}$ (E), and $Ni^{2+}$ (F)—show no significant fluorescence quenching, indicating the absence of complex formation. These data support the high selectivity of melatonin for $Cu^{2+}$ over biologically abundant metal ions.

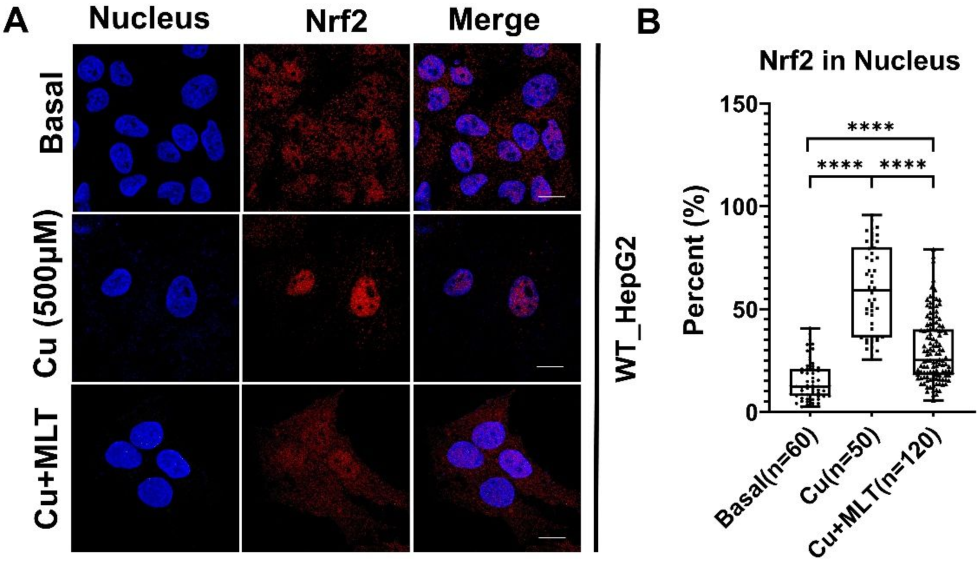


**Figure S5.**

**Nrf2 cytosolic relocalization in copper-exposed WT HepG2 cells after melatonin treatment**

(A) Representative immunofluorescence images showing Nrf2 localization in WT HepG2 cells under basal, copper-treated, and melatonin + copper conditions. Nuclei were counterstained with DAPI (blue); Nrf2 is shown in Red. (B) Quantification of nuclear Nrf2 localization in WT HepG2 cells reveals increased nuclear Nrf2 translocation upon copper exposure (~59.65% in WT), which was significantly reduced by melatonin pretreatment (~30% in WT) (mean ± SD, n=3, ***$p < 0.001$).

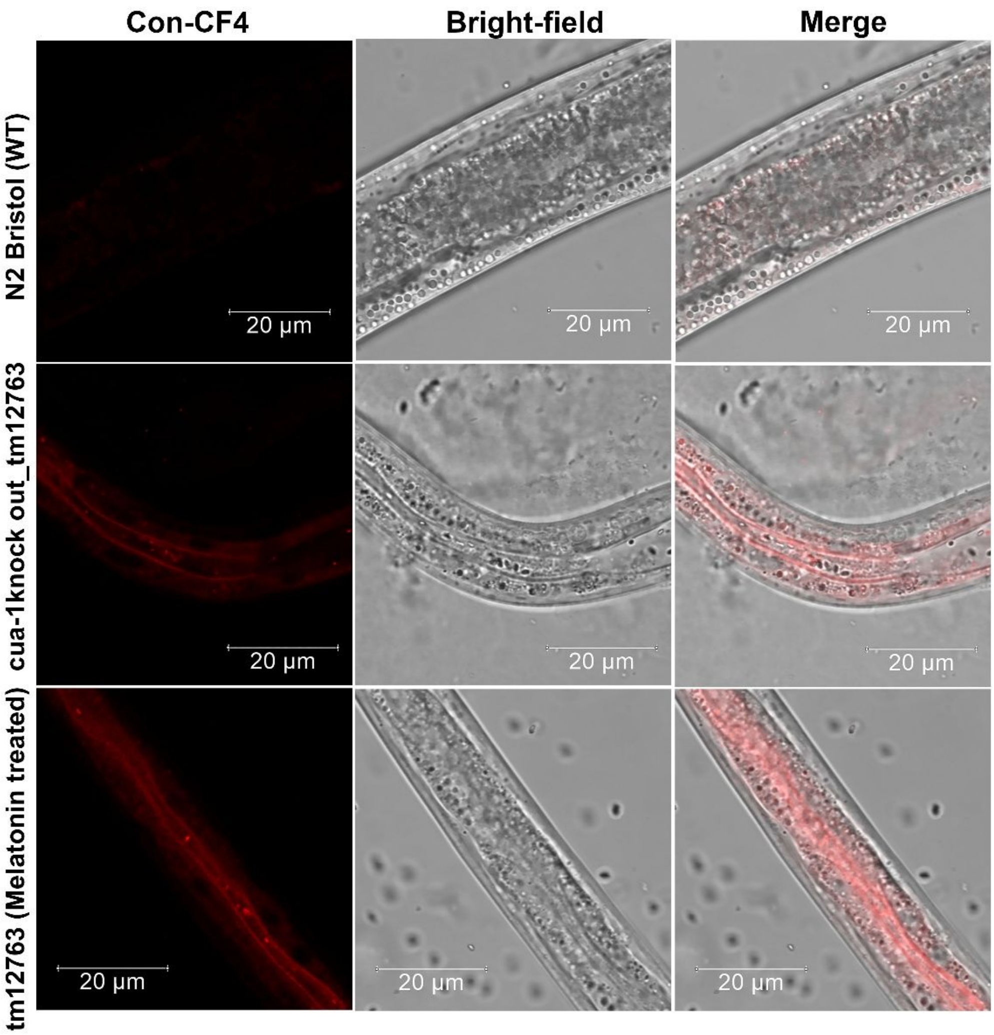


**Figure S6.**

**CF4 probe specificity validation**.
Wild-type and cua-1(tm12763) worms stained with control CF4 dye (non-responsive to copper) show minimal, diffuse fluorescence without a punctate signal, confirming the copper specificity of Cu-bound CF4. Scale bar: 20 μm.

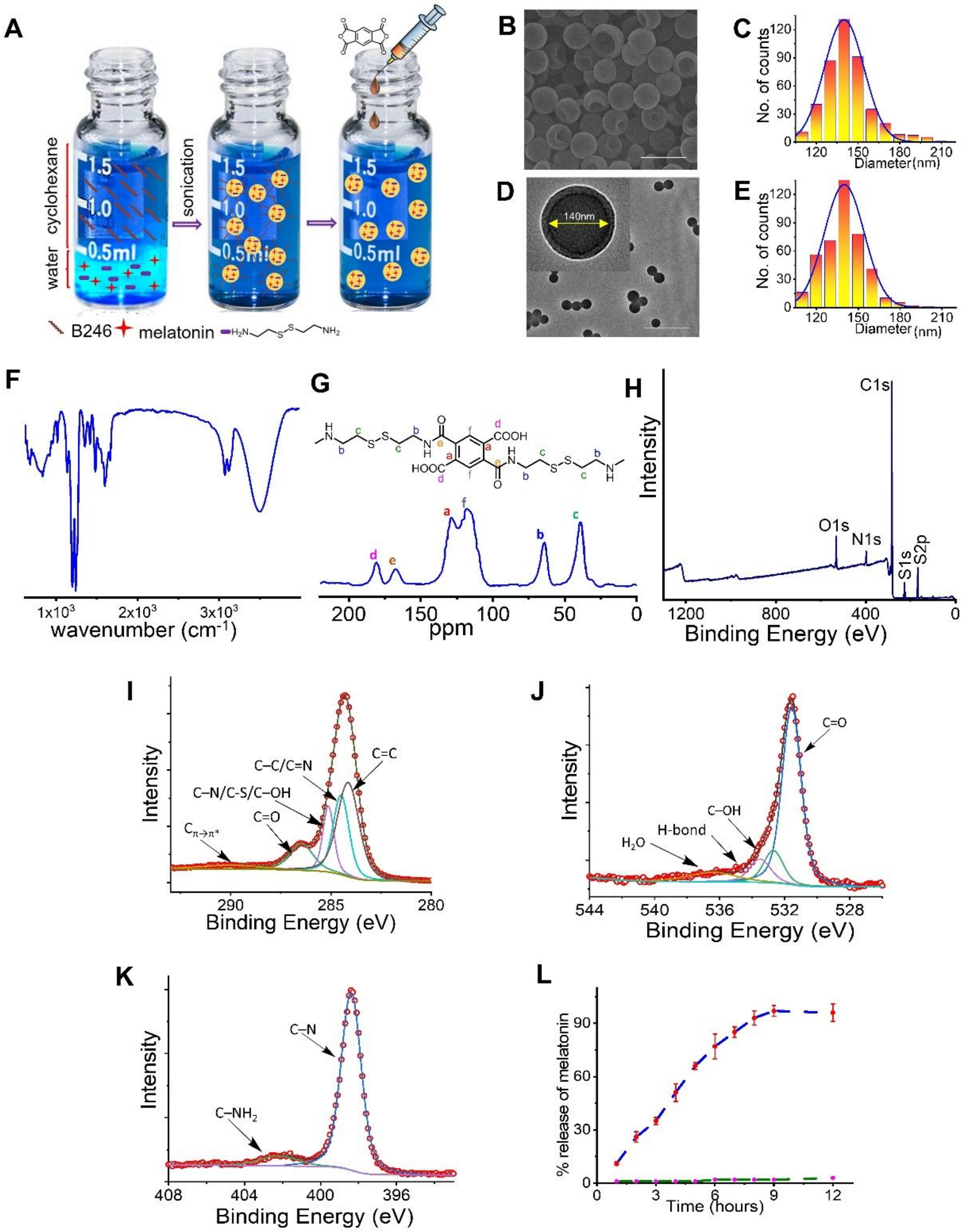


**Figure S7.**

**Synthesis and characterization of melatonin loaded nanocapsules**

(A). Illustration of synthesis of redox-responsive nanocapsules using an interfacial inverse mini-emulsion. (B-E). Morphological analysis of nanocapsules using SEM (B), DLS (C), and TEM (D), confirming spherical shape, narrow size distribution (140 ± 10 nm), and thin polymer shell (~16 ± 4 nm); (E) DLS reveals a mean hydrodynamic diameter of 160 nm with a PDI of 0.07. (F) FTIR spectrum confirming amide bond formation, with characteristic bands at 1643 $cm^{-1}$ (C=O stretch), 1571 $cm^{-1}$ (N–H bend), and 3495 $cm^{-1}$ (N–H/O–H stretch). (G) Solid-state $^{13}C$ NMR spectrum of lyophilized nanocapsules showing key peaks at 168 ppm (C=O amide), 181 ppm (−COOH), 110–140 ppm (aromatic carbons), 39 ppm (aliphatic C–S), and 63 ppm (aliphatic C–N). (H-K). XPS analysis confirming elemental composition and chemical environment: (H) survey spectrum identifying S 2p, S 1s, C 1s, N 1s, and O 1s; (I) deconvoluted C 1s showing C=C, C–C/C=N, C–N/C–S/C–OH, C=O, and π–π*; (J) O 1s peaks corresponding to C=O, C–OH, hydrogen bonding, and adsorbed water; (K) N 1s with peaks for C–N and free amine groups; (L) GSH-triggered melatonin release profile from nanocapsules at pH 7.4, showing ~95% release within 8 h under reducing conditions; negligible release observed in the absence of GSH.

**Legends to videos**

WT, ATP7B$^{-/-}$, and ATP7B$^{-/-}$ HepG2 cells treated with melatonin for 2 hours [ATP7B$^{-/-}$+MLT (2 hrs)] or 18 hours [ATP7B$^{-/-}$ +MLT (18 hrs)] were simultaneously excited using 405 nm and 488 nm lasers. The emission spectra were recorded between 500–550 nm under live conditions for 8 consecutive frames. Excitation with the 405 nm laser is shown in blue, while excitation with the 488 nm laser is shown in green. A merged video (blue + green) is also represented in each condition. Videos were acquired using a 63X oil immersion lens on the Leica SP8 confocal microscope. Scale bar- 20 μm

### Suppl material

**S3F**

**Chelation of Cu by melatonin**

In this study, we considered two probable pathways for the copper complexation with melatonin. These are, namely, the direct chelation mechanism (DCM) (equation 1) and the deprotonation-assisted chelation mechanism (DACM) (equation 2). The DCM pathway considers the complexation of melatonin (Mel) with free Cu cations, while the DACM involves the loss of amide/indole proton while complexing with Cu cations. Consideration of the DACM pathway is based on the observation from spectroscopic data, where loss of the amide proton peak was noticed.

$$Cu^{2+/+} + Mel \rightarrow [Mel - Cu]^{2+/+} \quad (1)$$

$$Cu^{2+/+} + Mel \rightarrow [Mel_{-H^+} - Cu]^{+/0} + H^+ \quad (2)$$

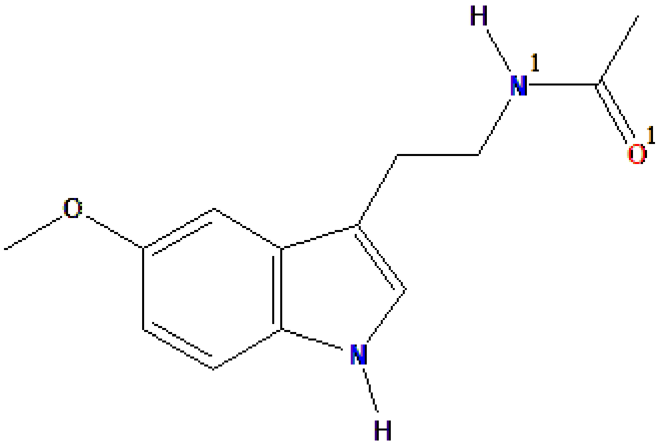


**Figure S3F-1**: Chemical structure of melatonin with coloring and numbering the chelation sites

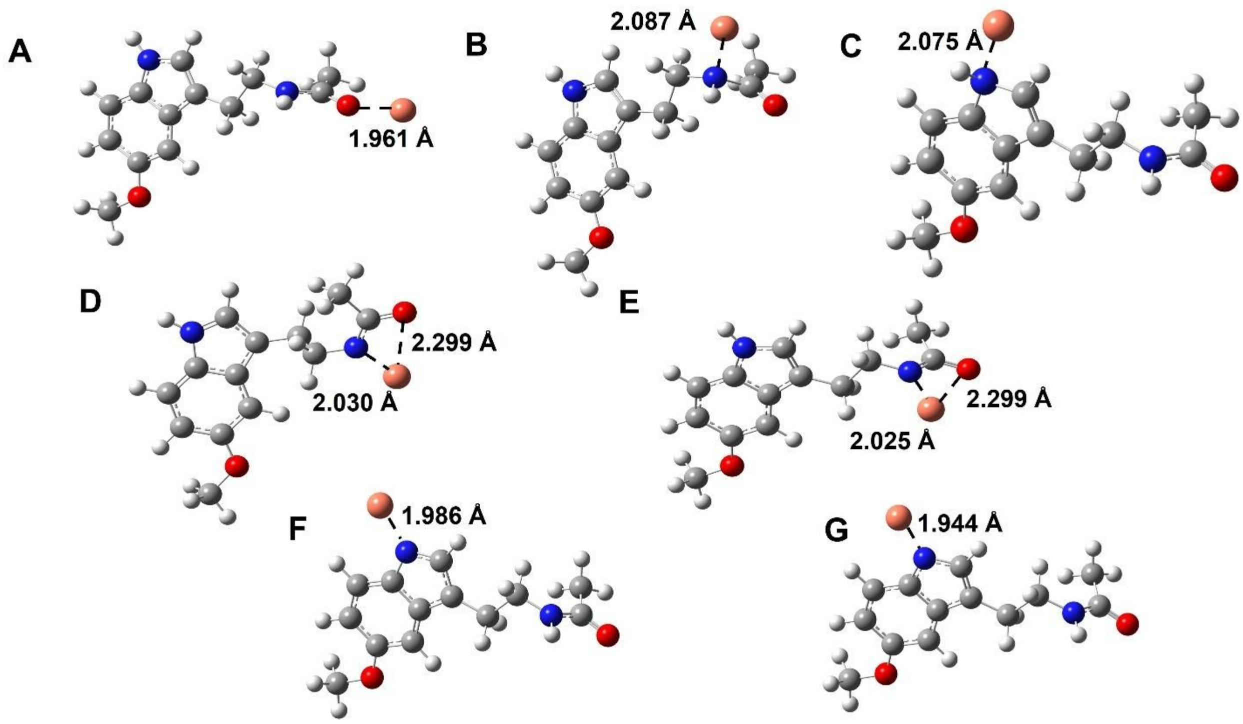


**Figure S3F-2:** Optimized geometries of melatonin–$Cu^{2+}$/$Cu^{+}$ complexes in aqueous medium, obtained at the M06-2X/6-311++G(d,p) level of theory. **A–C** show melatonin–$Cu^{+}$ complexes formed *via* the DCM through the amide O, amide N, and indole N, respectively. **D–G** represent complexes obtained through the DACM: D and E correspond to melatonin complexes with $Cu^{2+}$ and $Cu^{+}$ at the amide group, while F and G correspond to melatonin complexes with $Cu^{2+}$ and $Cu^{+}$ at the indole group. Distances between the potential chelation sites [O (*red*) and N (*blue*)] and the Cu atom (*brown*) are indicated with dotted lines.

**Table S3F-1:** Reaction free energies (ΔG) in kcal/mol for the melatonin and copper ion, and the atomic charge (in e units) the Cu atom in the corresponding complexes, *via* DCM and DACM pathways

| | | **Melatonin-$Cu^{2+}$** | | **Melatonin-$Cu^{+}$** | |
|---|---|---|---|---|---|
| | Chelation Site | ΔG | Atomic Charge on Cu | ΔG | Atomic Charge on Cu |
| **DCM** | Amide O | -95.76 | 0.95217 | -17.06 | 0.95089 |
| | Amide N | -86.56 | 0.95274 | -8.13 | 0.95063 |
| | Indole N | -74.64 | 0.99171 | -8.79 | 0.95171 |
| **DACM** | Amide N and O | 75.39 | 0.84553 | 155.09 | 0.83862 |
| | Indole N | 58.20 | 0.94093 | 152.67 | 0.88642 |

**Table S3F-2:** The donor-acceptor stabilization energy ($E^2$) in kcal/mol from second-order perturbation analysis of melatonin and Cu cation in melatonin—$Cu^{2+/+}$ complexes. Interaction energies exceeding 0.5 kcal/mol are only considered.

| | **Chelation Site** | **Donor** | **Acceptor** | **Melatonin-Cu (II)** | **Melatonin-Cu(I)** |
|---|---|---|---|---|---|
| **DCM** | Amide O | Amide O | Cu | 22.90 | 22.23 |
| | Amide N | Amide N | Cu | 15.34 | 15.85 |
| | Indole N | Indole N | Cu | 3.13 | 15.77 |
| **DACM** | Amide N and O | Amide O | Cu | 18.05 | 18.87 |
| | | Amide N | Cu | 34.35 | 35.74 |
| | Indole N | Indole N | Cu | 23.28 | 36.68 |

**S3G**

**RDF Calculation**

The radial distribution function (RDF) between two types of particles $a$ and $b$ is given by:

$$g(r) = \frac{1}{N_a N_b} \sum_{i=1}^{N_a} \sum_{j=1}^{N_b} \left\langle \delta\left(|\mathbf{r}_i - \mathbf{r}_j| - r\right)\right\rangle$$

which is normalized so that the RDF becomes 1 for large separations in a homogenous system. Here, $N_a$ and are the number of particles $a$ and $b$ in the system. $\delta$ is the Dirac delta function which counts the number of $b$ particles present around $a$ particle as distance $r$. In this case, $b$ is $Cu^{2+}$ ions and $a$ is either $N_1$, N or $O_1$.

**Table S1**

| Drug | Mechanism of action | Duration of therapy | Disadvantage |
|---|---|---|---|
| Dimercaprol | Copper chelation, | A few weeks or months | Painful intramuscular injections, Tachyphylaxis. |
| Penicillamine | Cu chelation | Lifelong | Worsening of neurological symptoms |
| Zinc salts | Decreases gastrointestinal Cu absorption | Lifelong | Biochemical pancreatitis, gastritis |
| Trientine | Cu chelation | Lifelong | Neurological deterioration, sideroblastic anemia |
